# Structural Basis of Protein Arginine Methyltransferase Activation by a Catalytically Dead Homolog (Prozyme)

**DOI:** 10.1101/776112

**Authors:** Hideharu Hashimoto, Lucie Kafková, Ashleigh Raczkowski, Kelsey Jordan, Laurie K. Read, Erik W. Debler

## Abstract

Prozymes are pseudoenzymes that stimulate the function of weakly active enzymes through complex formation. The major *Trypanosoma brucei* protein arginine methyltransferase, *Tb*PRMT1 enzyme (ENZ), requires *Tb*PRMT1 prozyme (PRO) to form an active heterotetrameric complex. Here we present the x-ray crystal structure of the ENZ-Δ52PRO tetrameric complex with the cofactor product S-adenosyl-L-homocysteine (AdoHcy) at 2.4 Å resolution. The individual ENZ and PRO units adopt the highly-conserved PRMT domain architecture and form an antiparallel heterodimer that corresponds to the canonical homodimer observed in all previously reported PRMTs. In turn, two such heterodimers assemble into a tetramer both in the crystal and in solution with twofold rotational symmetry. ENZ is unstable in absence of PRO and incapable of forming a homodimer due to a steric clash of a non-conserved tyrosine within the dimerization arm, rationalizing why PRO is required to complement ENZ to form a PRMT dimer that is necessary, but not sufficient for PRMT activity. The PRO structure deviates from other, active PRTMs in that it lacks the conserved η2 3_10_-helix within the Rossmann fold, abolishing co-factor binding. In addition to its chaperone function for ENZ, PRO substantially contributes to substrate binding. Heterotetramerization is required for catalysis, since heterodimeric ENZ-PRO mutants lack binding affinity and methyltransferase activity towards the substrate protein *Tb*RGG1. Together, we provide a structural basis for PRMT1 ENZ activation by PRO heterotetramer formation, which is conserved across all kinetoplastids, and describe a chaperone function of the PRMT1 prozyme, which represents a novel mode of PRMT regulation.

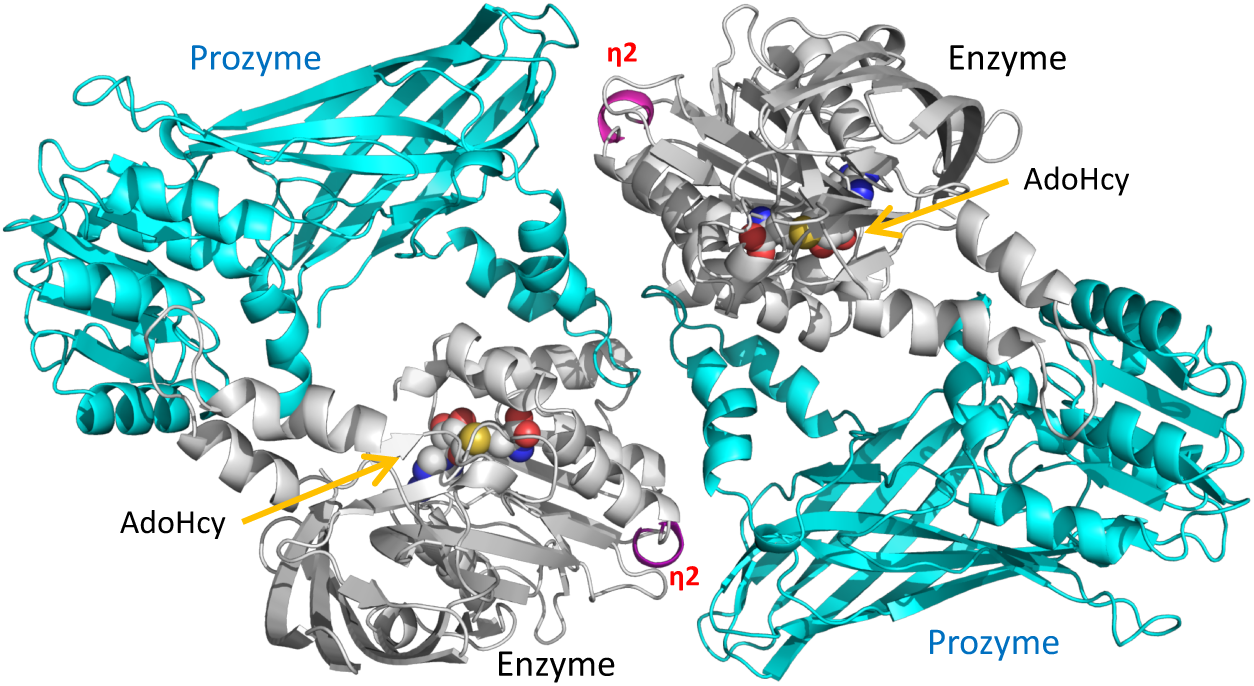
Graphical Abstract.

**Highlights:**

- The crystal structure of *Trypanosoma brucei* protein arginine methyltransferase 1 (PRMT1) is reported
- Two enzyme-prozyme heterodimers form a stable heterotetramer essential for catalysis
- The catalytically dead prozyme lacks elements essential for AdoMet binding
- The enzyme alone is unstable and cannot form a canonical dimer due to steric clashes
- *T. brucei* PRMT1 prozyme adopts a chaperone function conserved across kinetoplastids

## Introduction

*Trypanosoma brucei* is a protozoan parasite that can cause fatal sleeping sickness in humans and various animal diseases in Sub-Saharan Africa. An estimated 70 million people are at risk of infection [1]. African trypanosomes adopt two distinct replicative forms during their life cycle, the bloodstream form in the mammal and the procyclic form in the midgut of its insect vector, the tse-tse fly [2]. Control of gene expression and life-cycle progression primarily takes place on the post-transcriptional level [3]. However, recent evidence points towards a role of chromatin proteins in this process as well [4]. Among the posttranslational modifications that impact gene expression and other cellular functions, arginine methylation is prevalent with at least 15% of the proteome being modified, including proteins involved in a wide spectrum of processes such as RNA processing, DNA repair, metabolism, and protein trafficking [5, 6]. Arginine methylation is catalyzed by S-adenosyl-L-methionine (AdoMet)-dependent <u>p</u>rotein arginine (<u>R</u>) <u>m</u>ethyltransferases (PRMTs) [7]. Type I PRMTs (PRMT1, 2, 3, 4, 6 and 8) generate monomethyl arginine (MMA) and asymmetric dimethyl arginine (ADMA); type II PRMTs (PRMT5 and 9) generate monomethyl arginine (MMA) and symmetric dimethyl arginine (SDMA); and type III PRMTs (PRMT7) only produce monomethyl arginine [8, 9]. Their product specificities are restricted by the size and architecture of their active-site pockets [9–11].

*Tb*PRMT1 enzyme is the predominant type I PRMT1 in *T. brucei* that produces MMA and ADMA [12–14]. It contributes to parasite virulence, metabolic regulation, and nutritional stress response [15]. RGG/RG motifs are often the target of asymmetric arginine dimethylation in substrates, including the *Tb*PRMT1 substrates *Tb*RBP16 and *Tb*RGG1 [16]. The *Tb*PRMT1 enzyme shares 51% identical residues with type I human and rat PRMT1 [13, 17], and 41% identical residues with type I rat PRMT3 [18]. The PRMT structures suggest that the PRMT fold and the catalytic mechanism are conserved and that at least dimerization of the PRMT cores is required for AdoMet binding and catalysis [17–19]. Human, rat, and yeast PRMT1s are homo-oligomeric complexes with molecular weights of 300-400 kDa in solution [19–21], while rat PRMT3 exists in a monomer-dimer equilibrium in the cell with an activity of 0.3% with respect to rat PRMT1 [18, 22]. By contrast, *Tb*PRMT1 enzyme (ENZ) forms a stable heterotetrameric complex with a catalytically inert *Tb*PRMT1 prozyme (PRO), which was previously termed *Tb*PRMT3 [14]. The methyltransferase activity of ENZ by itself is not detectable, but catalytically inert PRO is necessary and sufficient to enable *Tb*PRMT1 activity. The protein expression levels of ENZ and PRO are strongly synchronized and interdependent [23], and we proposed that the amount of catalytic active *Tb*PRMT1 ENZ is regulated by the catalytic inert *Tb*PRMT1 PRO [14].

In addition to PRO, two other prozymes have been discovered in *Trypanosoma brucei* to date: AdoMet decarboxylase (AdoMetDC) and deoxyhypusine synthase (DHS) in the polyamine synthesis pathway [14, 24–27]. Prozymes are a subgroup of pseudoenzymes, which are estimated to represent ∼10% of the human proteome [28, 29] and are thought to serve as regulators of enzymes [25]. For example, the catalytically inert AdoMetDC prozyme facilitates enzyme catalysis by heterodimerization with the enzyme, and regulates enzymatic activity by dynamic allosteric changes at the active site [25].

ENZ and catalytically inactive PRO share several conserved motifs, including the Rossmann fold and motifs I and II (highlighted in bold in **Fig. 1**) with lower sequence identity (27%, 82 residues of 304), and similarity (44%, 136 residues of 304) [30]. A striking difference refers to conserved AdoMet binding residues [31, 32], many of which are lacking in PRO (**Fig. 1**). AdoMet crosslink experiments demonstrated that ENZ but not PRO binds AdoMet, consistent with the notion that PRO is a catalytically inactive enzyme [14].

**Figure 1.**
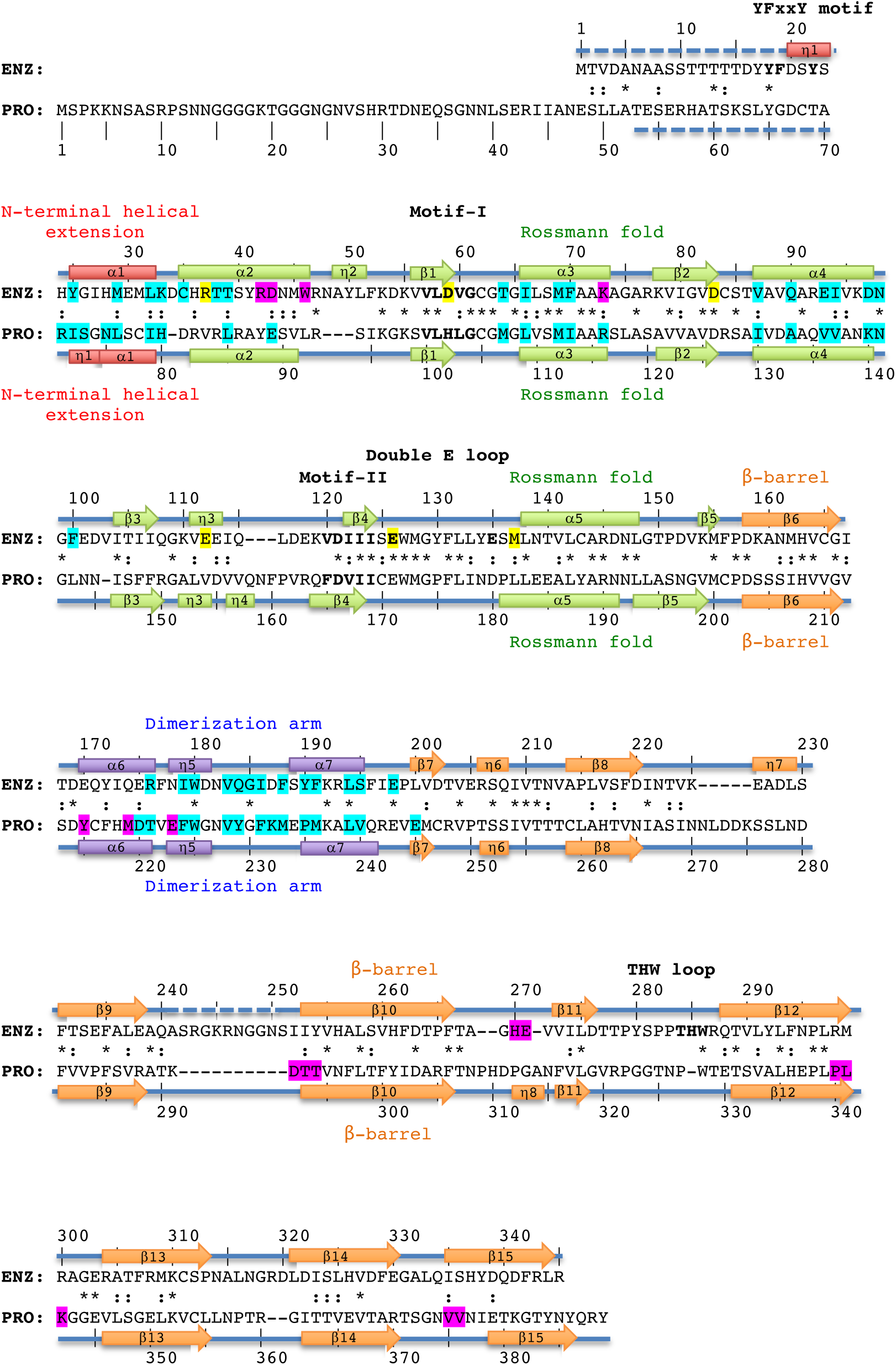
Structure-based sequence alignment of *Tb*PRMT1 enzyme and prozyme. Structure-based alignment of *Tb*PRMT1 enzyme (ENZ) and *Tb*PRMT1 prozyme (PRO). α1-α7 refer to *α*-helices, η1-η8 to 3_10_-helices, and β1-β15 to *β*-strands, indicating the secondary structure elements of *Tb*PRMT1 ENZ and PRO. Residue numbering is shown above (ENZ) and below (PRO) the sequences. Red designates the N-terminal helical extension (ENZ residues 20-33, PRO residues 71-80), green the Rossmann fold (ENZ residues 34-157, and PRO residues 81-202), orange the β-barrel (ENZ residues 158-345, and PRO residues 203-389), and purple the dimerization arm (ENZ residues 168-199, and PRO residues 213-244). Similar and identical residues are marked as : and *, respectively. Residues highlighted in magenta are involved in tetramer formation, residues highlighted in cyan are involved in ENZ-PRO dimer formation. Signature residues of the YFxxY motif, Motif-I, -II, the double E-loop, and the THW-loop are shown in bold. Key AdoMet-binding residues of ENZ are highlighted in yellow. Disordered regions are represented by dashed lines.

In order to elucidate the structural basis of enzyme activation by prozyme in *Tb*PRMT1, we determined the crystal structure of the ENZ-PRO complex, analyzed its oligomeric state in solution, and performed substrate binding and methyltransferase assays of wild-type and mutant ENZ and PRO species. Our results reveal that PRO is required for ENZ stability, that the heterotetrameric architecture of the ENZ-PRO complex is necessary for substrate binding and catalytic activity, and that the features of the *Tb*PRMT1 PRO-ENZ complex are conserved among kinetoplastids, implying a similar mode of PRMT1 regulation in these organisms.

## Results

### *Tb*PRMT1 ENZ-PRO forms a heterotetrameric complex

*Tb*PRMT1 ENZ-PRO from procyclic-form cells as well as the recombinantly expressed complex in *Escherichia coli* forms a heterotetramer as deduced from ultracentrifugation, size exclusion chromatography, and multiangle light scattering coupled to size exclusion chromatography (SEC-MALS) [14]. Because wild-type ENZ was insoluble or unstable [14] (**Table 1**), the wild-type ENZ (residues 1-345) and hexa-histidine-tagged wild-type PRO (residues 1-389) were co-expressed in *E. coli* using a pETDuet vector system (Novagen) and purified from the soluble fraction, followed by His-tag removal. According to SEC-MALS, the size of this complex was about 164 kDa, which corresponds to a heterotetrameric complex (theoretical molecular weight of 163.2 kDa), as previously described (**Table 1**) [14]. The heterotetrameric complex was further confirmed by small angle x-ray scattering coupled to SEC (SEC-SAXS) (**Fig. 2a**) and by negative-stain electron microscopy (EM) (**Fig. 2b**). In detail, the SEC-SAXS profile yielded a single peak with a constant radius of gyration across the peak (∼43.0 Å) (**Fig. 2a**). Based on the volume of correlation (V_c_) of this peak [33], a molecular weight of 187 kDa was estimated (**Table 2**). The Guinier and Kratky plots revealed that the protein was monodisperse and fully folded (**Fig. 2a**). The *Tb*PRMT1 ENZ-PRO complex visualized by negative-stain EM showed four distinct globular masses symmetrically arranged at the vertices of a rhomboid structure in the most predominant 2D class average (**Fig. 2b**). The complex measures about 140 Å in its longest dimension, which fits the maximum distance of 143 Å obtained in a pair-distance distribution by SEC-SAXS (**Fig. 2a**). Finally, a stoichiometry of 1:1 was obtained for the *Tb*PRMT1 heterotetramer with a Maltose Binding Protein (MBP)-fused *Tb*RGG1 substrate by isothermal titration calorimetry (ITC) (**Fig. 2c** and **Table 1**) [14]. Collectively, these data demonstrate that the *Tb*PRMT1 complex forms a rigid heterotetrameric unit in solution.

**Table 1.**
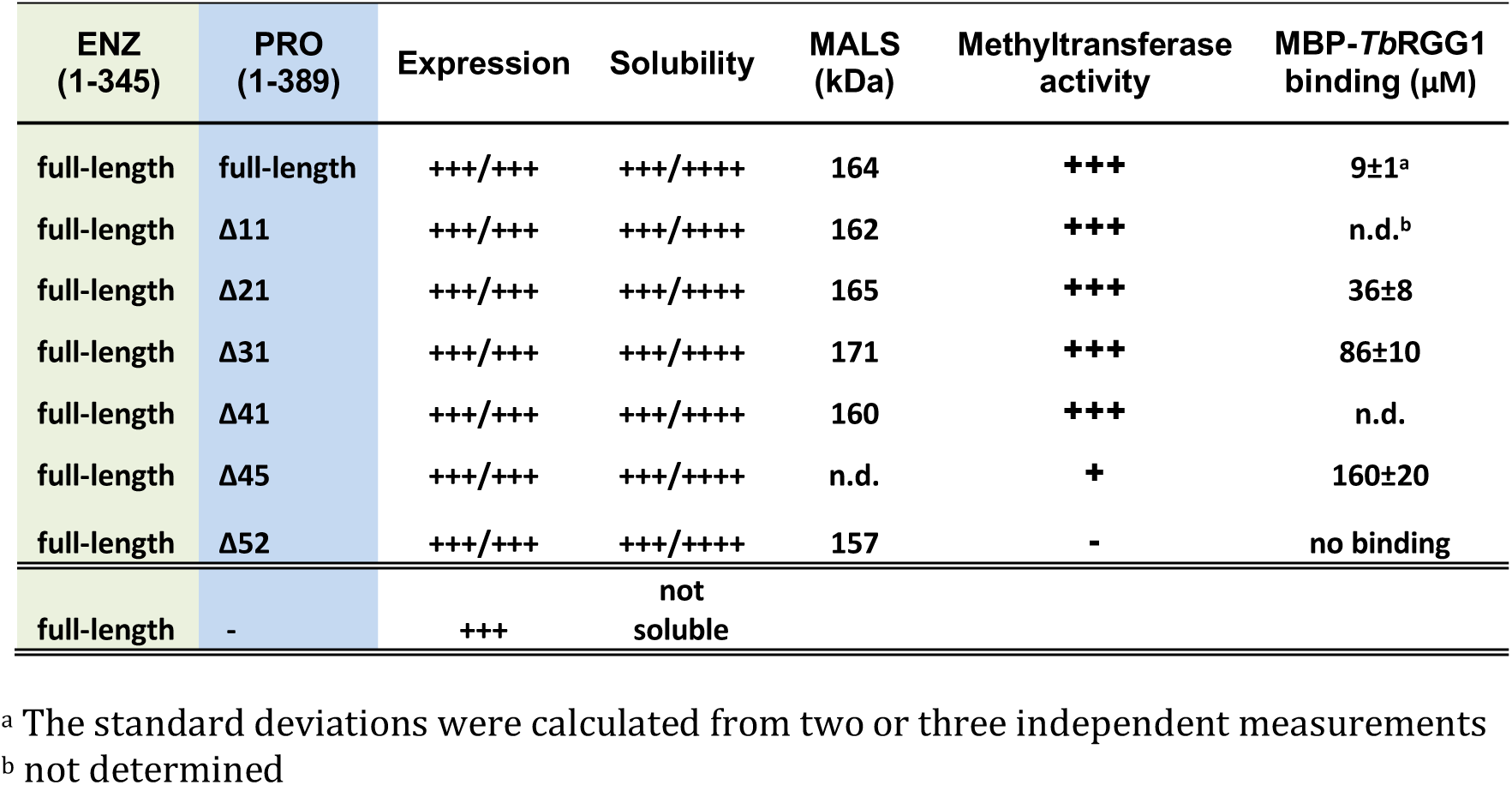
Methyltransferase activities, molecular weight, and binding affinities of ENZ-PRO and truncated PRO mutants.

**Table 2.**
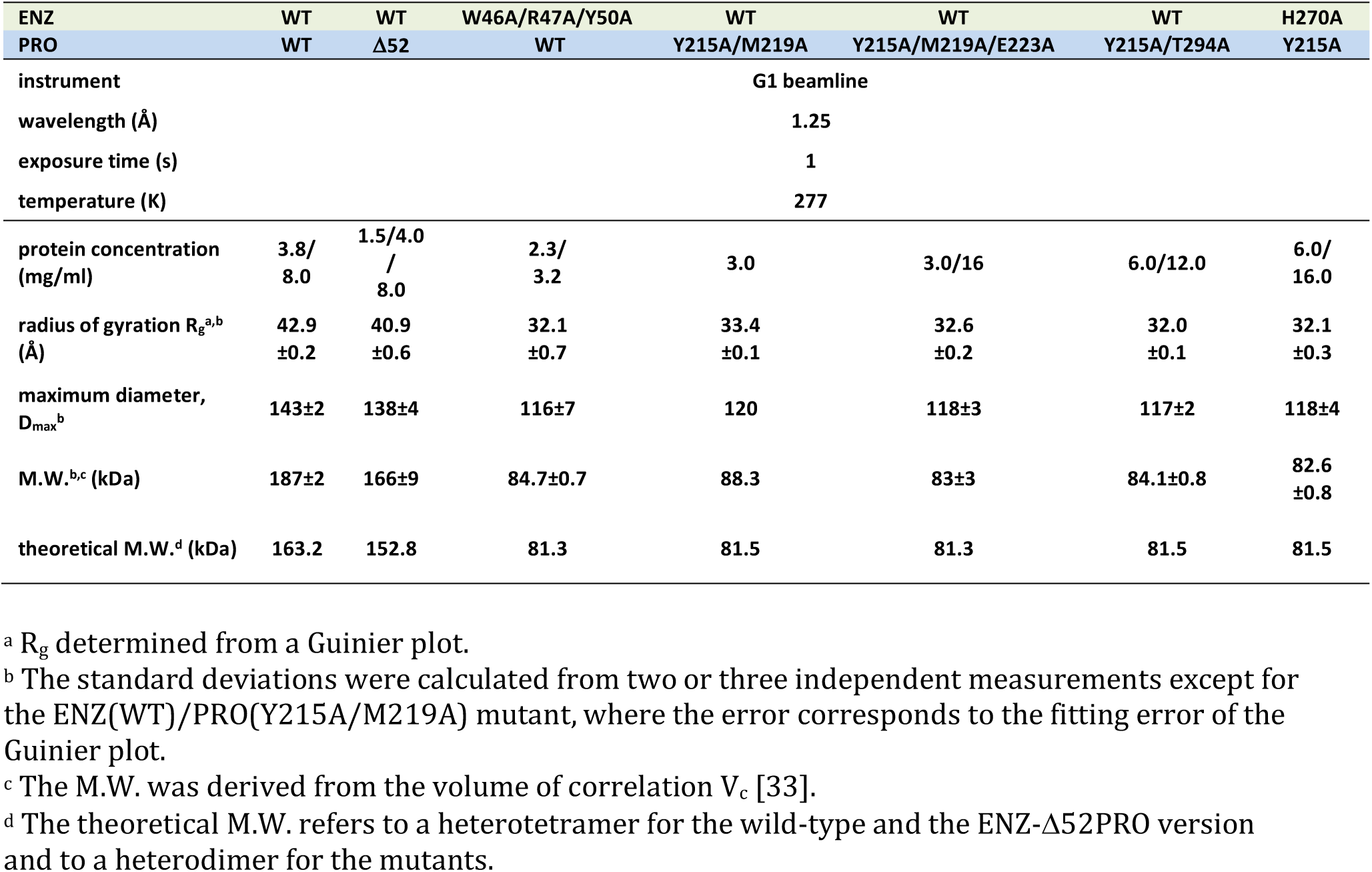
SEC-SAXS analysis of ENZ-PRO wild-type, ENZ-Δ52PRO, and interface mutants.

| ENZ | WT | WT | W46A/R47A/Y50A | WT | WT | WT | H270A |
| --- | --- | --- | --- | --- | --- | --- | --- |
| PRO | WT | Δ52 | WT | Y215A/M219A | Y215A/M219A/E223A | Y215A/T294A | Y215A |
| instrument | G1 beamline |  |  |  |  |  |  |
| wavelength (Å) | 1.25 |  |  |  |  |  |  |
| exposure time (s) | 1 |  |  |  |  |  |  |
| temperature (K) | 277 |  |  |  |  |  |  |
| protein concentration (mg/ml) | 3.8/<br>8.0 | 1.5/4.0<br>/<br>8.0 | 2.3/<br>3.2 | 3.0 | 3.0/16 | 6.0/12.0 | 6.0/<br>16.0 |
| radius of gyration $R_g^{a,b}$ (Å) | 42.9<br>±0.2 | 40.9<br>±0.6 | 32.1<br>±0.7 | 33.4<br>±0.1 | 32.6<br>±0.2 | 32.0<br>±0.1 | 32.1<br>±0.3 |
| maximum diameter, $D_{max}^b$ | 143±2 | 138±4 | 116±7 | 120 | 118±3 | 117±2 | 118±4 |
| M.W.<sup>b,c</sup> (kDa) | 187±2 | 166±9 | 84.7±0.7 | 88.3 | 83±3 | 84.1±0.8 | 82.6<br>±0.8 |
| theoretical M.W.<sup>d</sup> (kDa) | 163.2 | 152.8 | 81.3 | 81.5 | 81.3 | 81.5 | 81.5 |

<sup>a</sup> $R_g$ determined from a Guinier plot.
<sup>b</sup> The standard deviations were calculated from two or three independent measurements except for the ENZ(WT)/PRO(Y215A/M219A) mutant, where the error corresponds to the fitting error of the Guinier plot.
<sup>c</sup> The M.W. was derived from the volume of correlation $V_c$ [33].
<sup>d</sup> The theoretical M.W. refers to a heterotetramer for the wild-type and the ENZ-Δ52PRO version and to a heterodimer for the mutants.

**Figure 2.**
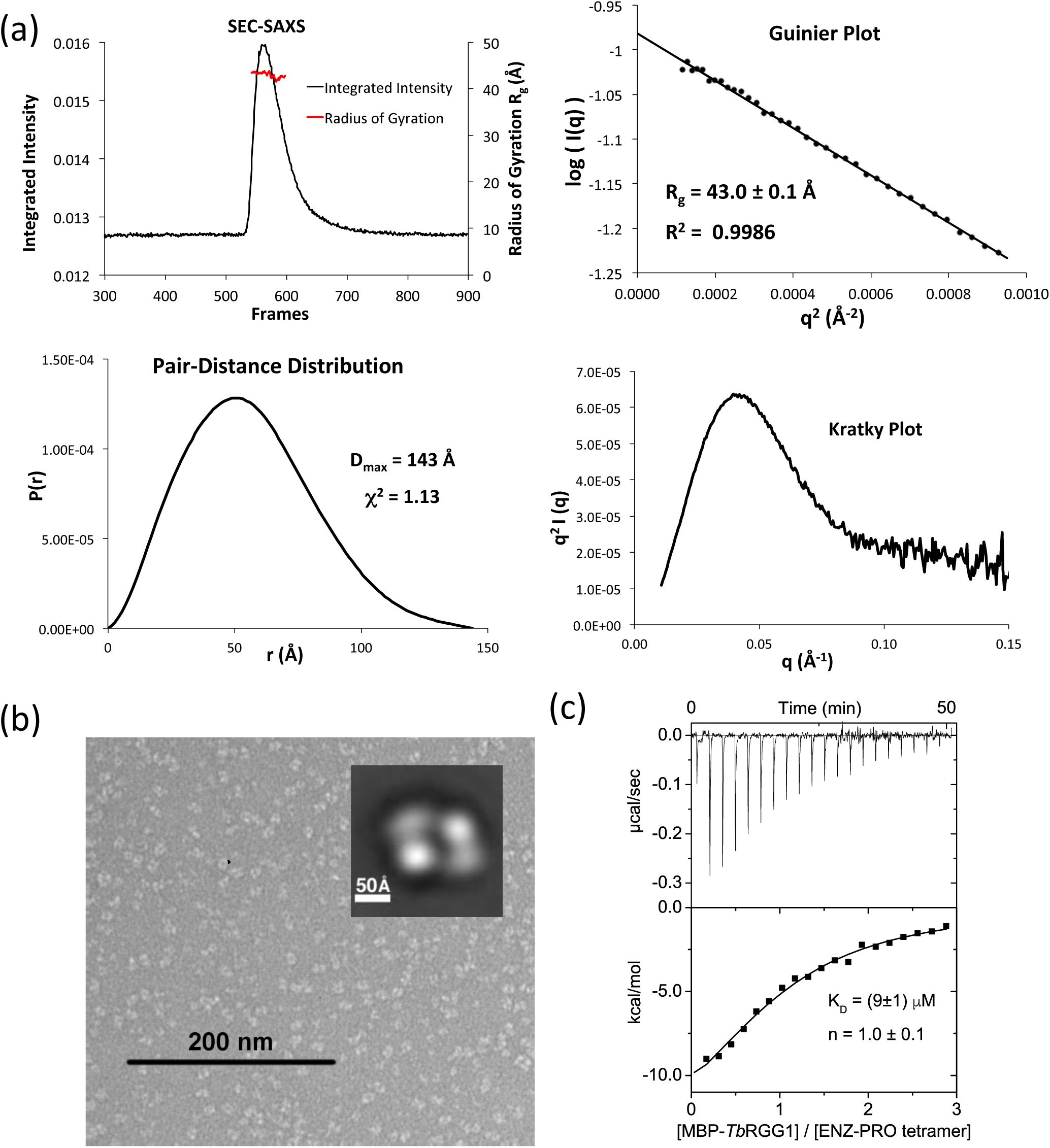
*Tb*PRMT1 ENZ-PRO forms a heterotetrameric complex in solution. (a) SEC–SAXS analysis of *Tb*PRMT1 ENZ-PRO. Top left panel: SEC-SAXS integrated intensities (left y-axis) plotted against frame number (x-axis). The red dots indicate radius of gyration, R_g_ (on the right y-axis). Top right panel: Guinier plot calculated from averaging buffer-subtracted scattering intensities. The coefficient of determination, R^2^, is 0.9986. Bottom left panel: Pair-distance distribution function P(r), yielding a maximum molecular diameter of 143 Å. Bottom right panel: Normalized Kratky plot calculated from SEC–SAXS data. (b) Negative-stain electron microsopy. EM micrograph with a 200 nm scale bar. Inset: Predominant 2D class average. (c) ITC thermogram (upper panel) and plots of corrected heat values (lower panel) for binding of the heterotetrameric ENZ-PRO complex to Maltose Binding Domain (MBP)-fused *Tb*RGG1 protein.

### The *Tb*PRMT1 PRO N-terminus contributes to substrate recognition

Previous studies have shown that all PRMTs contain a highly conserved core domain comprising ∼310 residues [8] and that the N-terminal region of PRMTs is often flexible and involved in substrate recognition [34, 35]. Therefore, we examined the role of the N-terminal residues of PRO and ENZ. Limited proteolysis on the full-length proteins identified a stable, N-terminally truncated PRO fragment spanning residues 53-389, referred to as Δ52PRO, while ENZ remained intact under the conditions tested. The Δ52PRO fragment was then co-expressed with full-length ENZ. SEC-MALS, SEC-SAXS, and SEC alone confirmed that the ENZ-Δ52PRO complex still formed a tetrameric complex (**Fig. S1a** and **Tables 1** and **2**). However, its binding affinity for MBP-*Tb*RGG1 was abolished (**Fig. S1b)**, consistent with methyltransferase inactivity (**Table 1**). These data suggest that the N-terminus of PRO is essential for substrate binding. When we measured the methyltransferase activities and MBP-*Tb*RGG1 protein binding affinities of a series of N-terminal PRO deletion mutants, we found that the first ∼40 amino-terminal residues of PRO are dispensable for methyltransferase activity (**Table 1**). These data imply that the flexible N-terminal region between residues 41 and 52 of PRO is critical for substrate recognition.

### ENZ and PRO form heterodimers that assemble into a heterotetramer

In order to obtain mechanistic insights into ENZ activation by PRO at atomic resolution, we crystallized the stable ENZ-Δ52PRO complex with the methylation cofactor product AdoHcy and solved the structure at 2.4 Å resolution from a seleno-methionine derivatized crystal using the single anomalous dispersion (SAD) phasing technique (**Supplemental Table S1**). Two ENZ (shown in gray) and two Δ52PRO (shown in cyan) molecules form the asymmetric unit (**Fig. 3a**). No electron density of the ENZ residues 1-20, the ENZ loop region between β9-β10 strands (residues 241-250), and Δ52PRO residues 53-70 was observed. Consistent with our previous biochemical finding [14], AdoHcy was only bound to ENZ (**Fig. 3a**). Notably, one ENZ and one Δ52PRO molecule form a canonical PRMT ring-like dimeric structure that has been observed in all other, homodimeric PRMTs thus far [17–19, 36] (**Fig. 3b**). In turn, two ENZ-Δ52PRO heterodimers touch each other side by side, exhibiting a 2-fold non-crystallographic symmetry (**Fig. 3a**). The two dimers are highly similar with a root-mean-square deviation (rmsd) of only 0.6 Å when comparing 637 pairs of Cα atoms. The dimensions of the *Tb*PRMT1 ENZ-Δ52PRO heterotetrameric complex are 131 Å x 70 Å x 78 Å (**Fig. 3a**). Importantly, the molecular size and shape of the crystal structure are in good agreement with the results of SEC-MALS (**Table 1**), SEC-SAXS (**Supplemental Figs. S1a, Fig. 2a**, and **Table 2**), and electron microscopy (**Fig. 2b**), therefore corroborating the tetrameric form in solution, which is further confirmed by comparing the calculated radii of gyration of the ENZ-Δ52PRO heterotetramer and heterodimer (39 Å and 28 Å, respectively) with the experimental radius of gyration (41Å) [37].

**Figure 3.**
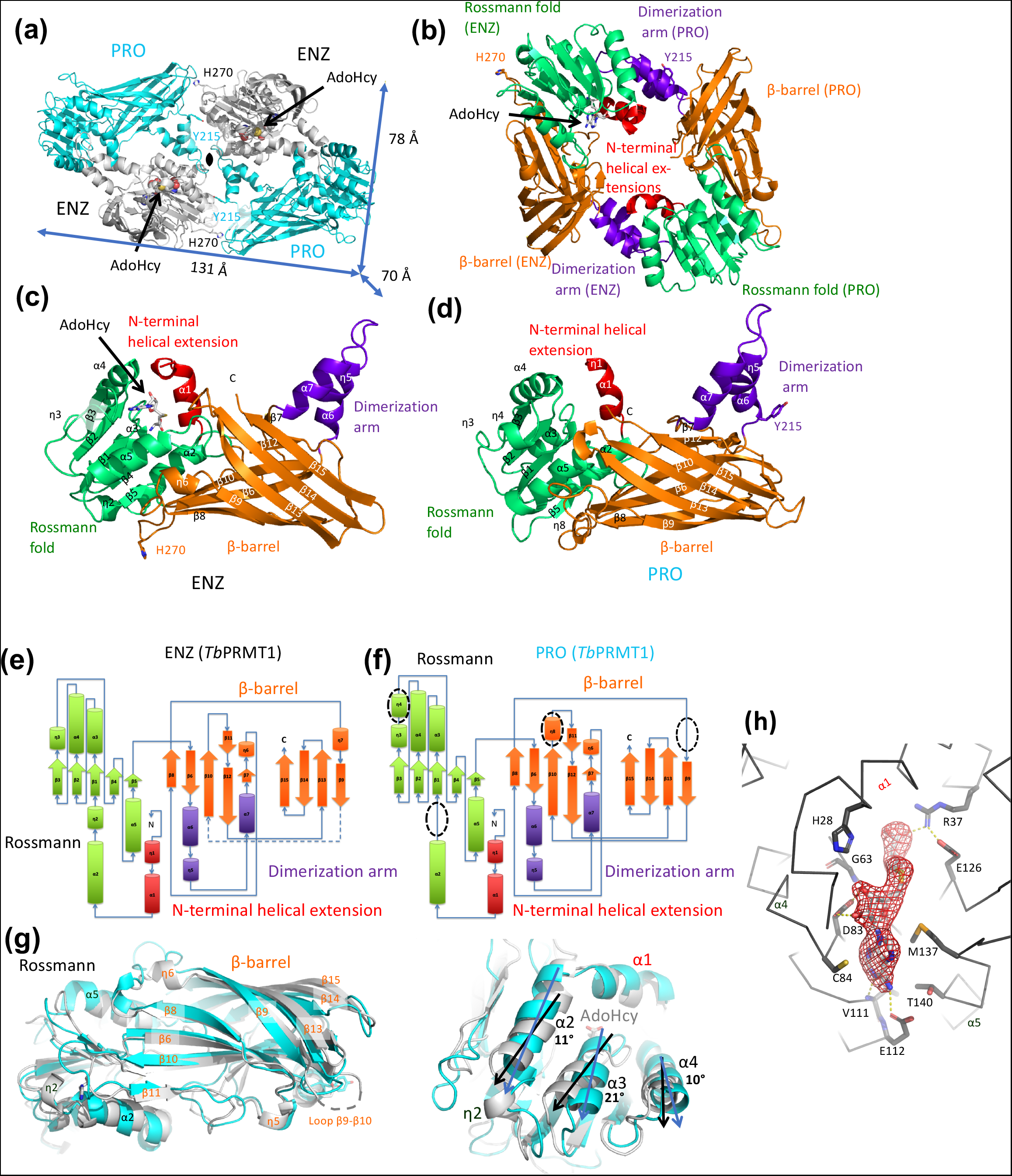
Crystal structure of heterotetrameric *Tb*PRMT1 ENZ-PRO complex. (a) Tetrameric *Tb*PRMT1 ENZ-PRO complex. One ENZ (gray) bound to AdoHcy shown in space-filling representation forms a heterodimer with PRO (cyan). The dimensions of the tetrameric complex are shown. The non-crystallographic twofold symmetry axis is shown as a black pointed oval. Key residues involving tetramerization (His270 of ENZ, and Tyr215 of PRO) are indicated. (b) Anti-parallel *Tb*PRMT1 ENZ-PRO heterodimer. The N-terminal helical extension (red), the Rossmann fold (green), the β-barrel (orange) and the dimerization arm (purple) are shown. AdoHcy, His270, and Tyr215 are highlighted. (c) Monomeric *Tb*PRMT1 ENZ structure. (d) Monomeric *Tb*PRMT1 PRO structure. (e) Topology of *Tb*PRMT1 ENZ. (f) Topology of *Tb*PRMT1 PRO. (g) Superimposition of monomeric ENZ and PRO. Left panel: Overall structure. Right panel: Close-up view on Rossmann fold, illustrating rotated *α*-helices. (h) Simulated-annealing omit electron density map for AdoHcy, contoured at 3.5*σ* above the mean.

### ENZ adopts the canonical PRMT fold and features a type I active-site architecture

ENZ harbors seven α-helices, six 3_10_-helices, and 15 β-strands (**Figs. 1, 3c**, and **3e**). The two ENZ molecules within the asymmetric unit are highly similar (rmsd of 0.4 Å, comparing 318 pairs of Cα atoms). The overall monomeric structure of ENZ strongly resembles the monomeric class I PRMT core domain structure of rat PRMT1, rat PRMT3, mouse CARM1 (PRMT4), and yeast RMT1 (**Supplemental Table S2**), sharing an identical topology with the rat PRMT1 core domain (**Fig. 3e** and **Supplemental Fig. S2**) [17–19, 36]. Like other PRMTs, ENZ contains the four highly conserved modules of PRMTs: an N-terminal helical extension (residues 20-33 in red), a Rossmann fold (residues 34-157 in green), a dimerization arm (residues 168-199 in purple), and a β-barrel domain (residues 158-345 in orange) (**Figs. 3b, 3c**, and **Supplemental Fig. S2**). The electron density of the co-factor product AdoHcy is clearly observed in both ENZ molecules (**Figs. 3a** and **3h**). AdoHcy interacts with residues that are highly conserved among human PRMT1, rat PRMT1, and *Tb*PRMT1 ENZ for AdoMet binding [17, 18, 31]: Tyr22, His28, Arg37, Asp83, Cys84, Glu112, Glu126, Met137, Thr140, the main chain of Gly63 and Val111, and Asp59 via two water molecules (**Fig. 3h**). The *Tb*PRMT1 ENZ active site possesses the previously described PRMT type I features [9–11]: An open subregion A adjacent to the double E-loop, while subregion B towards the conserved THW loop is sterically more restricted, which enables conversion to mono- and asymmetric dimethyl arginine, but not to symmetric dimethylarginine (**Supplemental Fig. S3**). The distances between atoms of conserved ENZ residues and of the sulfur atom of AdoHcy recapitulate the type I enzyme active-site architecture and provide a structural basis for *Tb*PRMT1 ENZ product specificity [14].

### Lack of the η2 3_10_-helix is a unique feature of the PRO core domain that twists its Rossmann fold

The two *Tb*PRMT1 Δ52PRO molecules (cyan in **Fig. 3a**) within the asymmetric unit superimpose closely (rmsd of 0.4 Å, comparing 319 pairs of Cα atoms) and harbor the four canonical PRMT modules: an N-terminal helical extension (residues 71-80 in red), a Rossmann fold (residues 81-202 in green), a dimerization arm (residues 213-244 in purple), and a β-barrel domain (residues 203-389 in orange) (**Figs. 3b** and **3d**). While the monomeric structures of ENZ and Δ52PRO are relatively similar (**Fig. 3g**, rmsd of 2.1 Å, comparing 293 pairs of Cα atoms), the topology of Δ52PRO bears a few marked differences with respect to ENZ and other PRMT1s (**Figs. 3e** and **3f**) [17–19, 36]. Within the Rossmann fold, PRO lacks the η2 3_10_-helix between the α2 helix and β1 strand, and has an extra η4 3_10_-helix between the η3 3_10_-helix and β4 strand (**Figs. 1** and **3f**). Importantly, the η2 3_10_-helix is highly conserved among active enzymes (**Supplemental Fig. S4**) [8]. As a result of the lacking η2 3_10_-helix, the α2, α3 and, α4 helices within the PRO Rossmann fold are tilted by 10-20°, whereas the β sheets of the ENZ and PRO Rossmann folds align well (**Fig. 3g**). In turn, these observed differences in the secondary structure elements of PRO twist and hence affect the dimerization interface, compromising cofactor AdoMet binding and providing a structural basis for PRO inactivity.

### PRO is required for dimerization and ENZ stability

The η1 3_10_-helix and the α1 and α2 helices contribute to AdoMet binding (**Fig. 3h**) and dimerization of PRMTs (**Fig. 3b**) [8]. The total buried dimerization surface area between ENZ and PRO is ∼1,600 Å^2^, which is similar to that of other, homodimeric PRMTs. Since dimerization arm mutants that lead to monomeric PRMTs do not have methyltransferase activity [10, 17, 19], dimerization of PRMTs is required for activity [9, 10, 17–19, 34, 36, 38]. As we previously reported, ENZ expressed by itself is an unstable protein that requires PRO to form a stable, catalytically active complex [14]. PRO, on the other hand, can be expressed by itself, albeit at a reduced amount, indicating decreased stability, and forms a homodimer [14]. Thus, the protein amount of folded ENZ is limited and hence may be regulated by PRO *in vivo* [14, 23]. As the molecular surface of the monomeric ENZ structure displays highly hydrophobic patches, covering these hydrophobic regions via dimerization with PRO is a likely mechanism to stabilize the ENZ protein. Particularly the dimerization arm of ENZ is highly hydrophobic and dominated by aromatic residues (**Fig. 4a**). In detail, the hydrophobic residues Ile179, Trp180, Val183, Ile186, Phe188, Tyr190, Phe191, and Leu194 of the ENZ dimerization arm contact the hydrophobic residues Ile72, Leu76, Ile79, Leu85, Met107, Leu109, Ile113, Ile130, Ala133, and Val137 on the η1 3_10_-helix, α1, α2, and α3 helices of PRO (**Fig. 4b**). Similarly, the surface of the PRO dimerization arm is hydrophobic. The PRO residues Thr221, Phe224, Trp225, Val228, Tyr229, Phe231, Met233, Pro235, Met236, Leu239, and Val240 contact the hydrophobic ENZ residues Tyr25, Met29, Lys33, Cys35, Thr38, Thr39, Arg42, Trp46, Thr64, Ile66, Phe70, Val87, Gln90, Ile94 and Phe100 on the η1 3_10_-helix, α1, α2, and α3 helices (**Fig. 4c**). Upon dimerization of ENZ and PRO, the hydrophobic patches of the η1 3_10_-helix and α1 helix in ENZ become buried, leading to ENZ stabilization.

**Figure 4.**
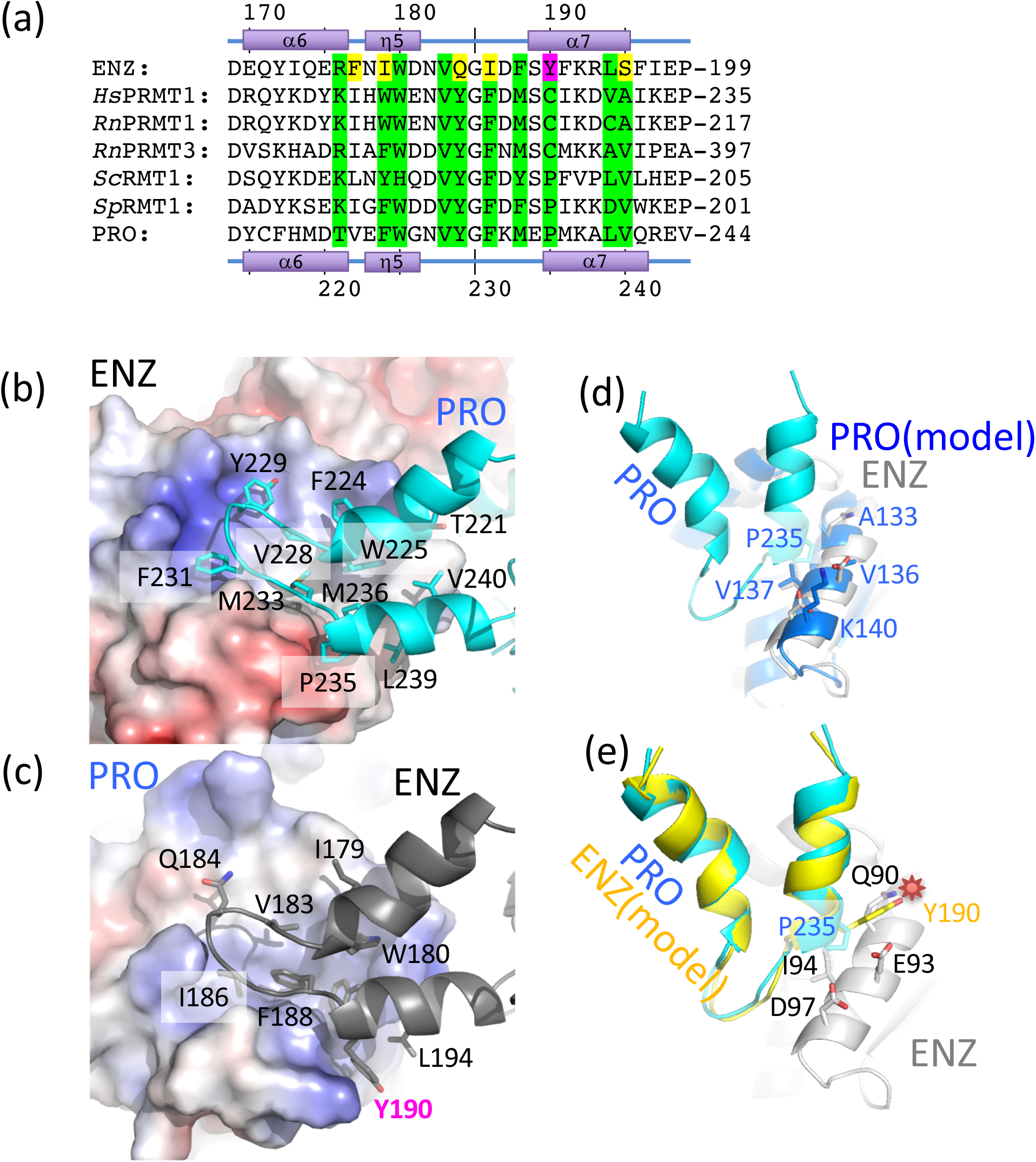
The dimerization arm of ENZ features non-conserved residues, preventing ENZ homodimerization. (a) Sequence alignment of the dimerization arm regions of *Tb*PRMT1 ENZ (TriTrypDB ID: **<u>Tb927.1.4690</u>**), rat PRMT1 (GenBank ID: **<u>NP_077339</u>**), rat PRMT3 (**<u>NP_4460009</u>**), budding yeast RMT1 (**<u>NP_009590</u>**), fission yeast RMT1 (**<u>NP_594825</u>**), and *Tb*PRMT1 PRO (TriTrypDB ID: **<u>Tb927.10.3560</u>**). Conserved interface residues are colored in green. Non-conserved interface residues are colored in yellow. The non-conserved ENZ Tyr190 is highlighted in magenta. (b) Hydrophobic interaction of the PRO dimerization arm with the ENZ Rossmann fold. (c) Hydrophobic interaction of the ENZ dimerization arm with the PRO Rossmann fold. (d) Modelled PRO-PRO interface. (e) Modelled ENZ-ENZ interface. The non-conserved Tyr190 causes a steric clash between two ENZ molecules highlighted by a red star, which interferes with dimerization.

The two dimerization arms of ENZ and PRO are structurally highly similar (rmsd of 0.7 Å, comparing 124 pairs of Cα atoms). When we generate homodimeric models of ENZ and PRO, respectively, we observe that PRO can indeed form homodimers without any severe steric clashes (**Fig. 4d**), while Tyr190 of ENZ, corresponding to Pro235 of PRO, clearly clashes with the side chains of Gln90, Glu93, and Ile94 on the α4 helix of ENZ (**Fig. 4e**). Consequently, ENZ is predicted to be unable to form a homodimer due to steric clashes and requires PRO to engage into a stable ENZ-PRO complex.

### The *Tb*PRMT1 heterotetrameric assembly is required for substrate binding and catalytic activity

Human and rat PRMT1 predominantly form oligomeric structures [17, 19–21], which are the active species *in vivo* [39]. PRMT3 exists in a monomer-dimer equilibrium in solution [18, 22]. However, PRMT3 crystal structures revealed the canonical dimeric arrangement, consistent with the notion that PRMT dimerization is necessary for catalytic activity [17–19]. *Tb*PRMT1 ENZ and PRO form a stable heterotetramer (**Figs. 2** and **3a**) [14] that binds one substrate molecule (**Fig. 2c**). The interface between two heterodimers amounts to ∼950 Å^2^ and is dominated by van der Waals’ contacts; Tyr50 of ENZ interacts with hydrophobic residues Val375 and Val376 of PRO, His270 of ENZ interacts with residues Asp292, Thr293, Thr294, Pro340, Leu341, and Val375 of PRO, Tyr215 of PRO interact with residues Asp43, Trp46, and Arg47 of ENZ, and Met219 of PRO interacts with Trp46 of PRO (**Fig. 5**). We mutated various key dimer-dimer surface residues to alanine in order to test whether they break down the tetramer into dimers, and evaluated the size of the resulting mutants using SEC, SEC-MALS, SEC-SAXS, and EM (**Fig. 6, Tables 2** and **3**). Indeed, the ENZ triple mutant W46A/R47A/Y50A, the ENZ double mutant Y50A/H270A, the PRO double mutants Y215A/M219A, Y215A/E232A, Y215A/K232A, Y215A/T294A, as well as the PRO triple mutant Y215A/M291A/E223A all disrupt the tetrameric complex and form a stable heterodimer.

**Table 3.**
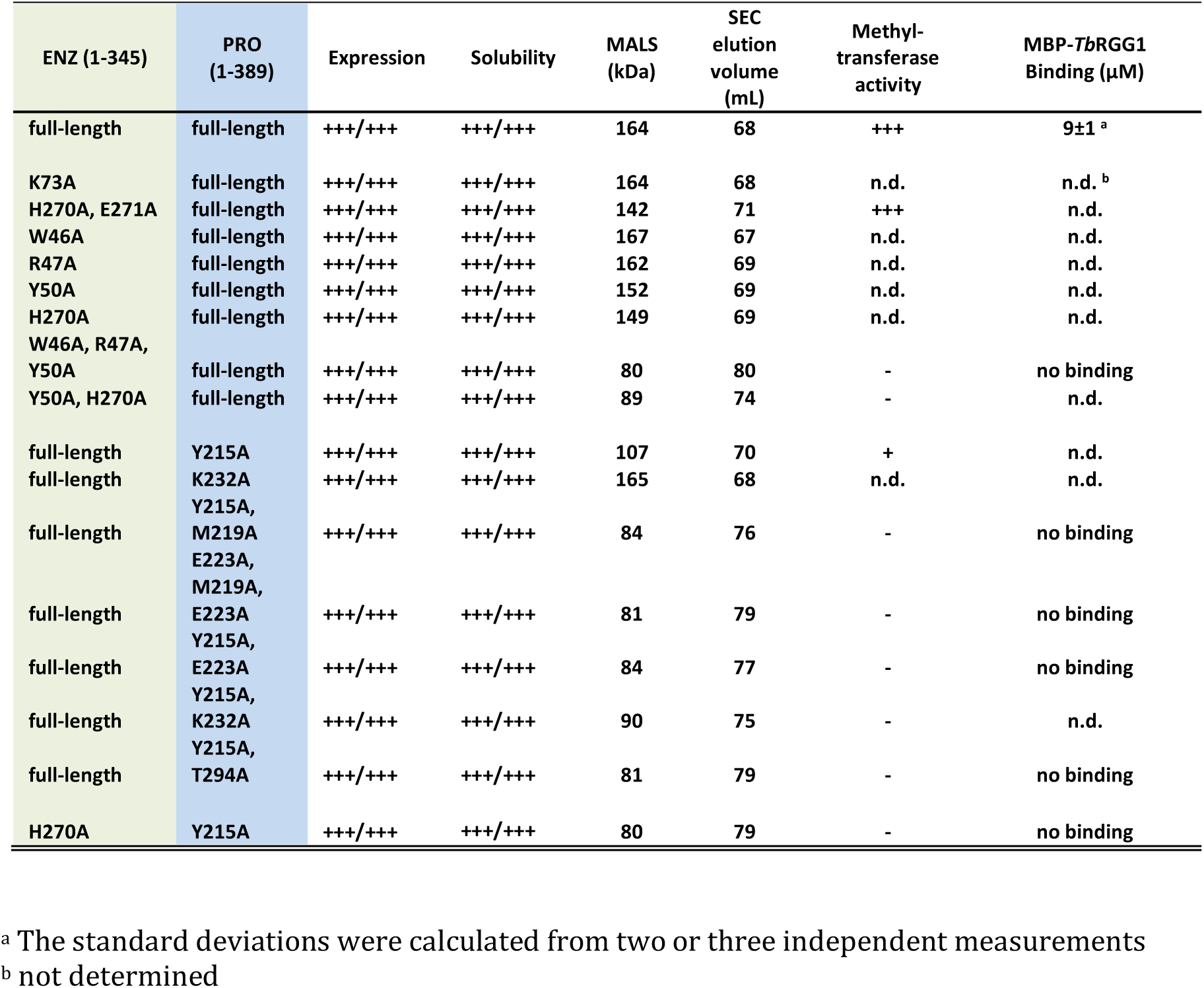
Methyltransferase activity, molecular weight, and MBP-*Tb*RGG1 binding affinities of ENZ-PRO and tetrameric interface mutants.

**Figure 5.**
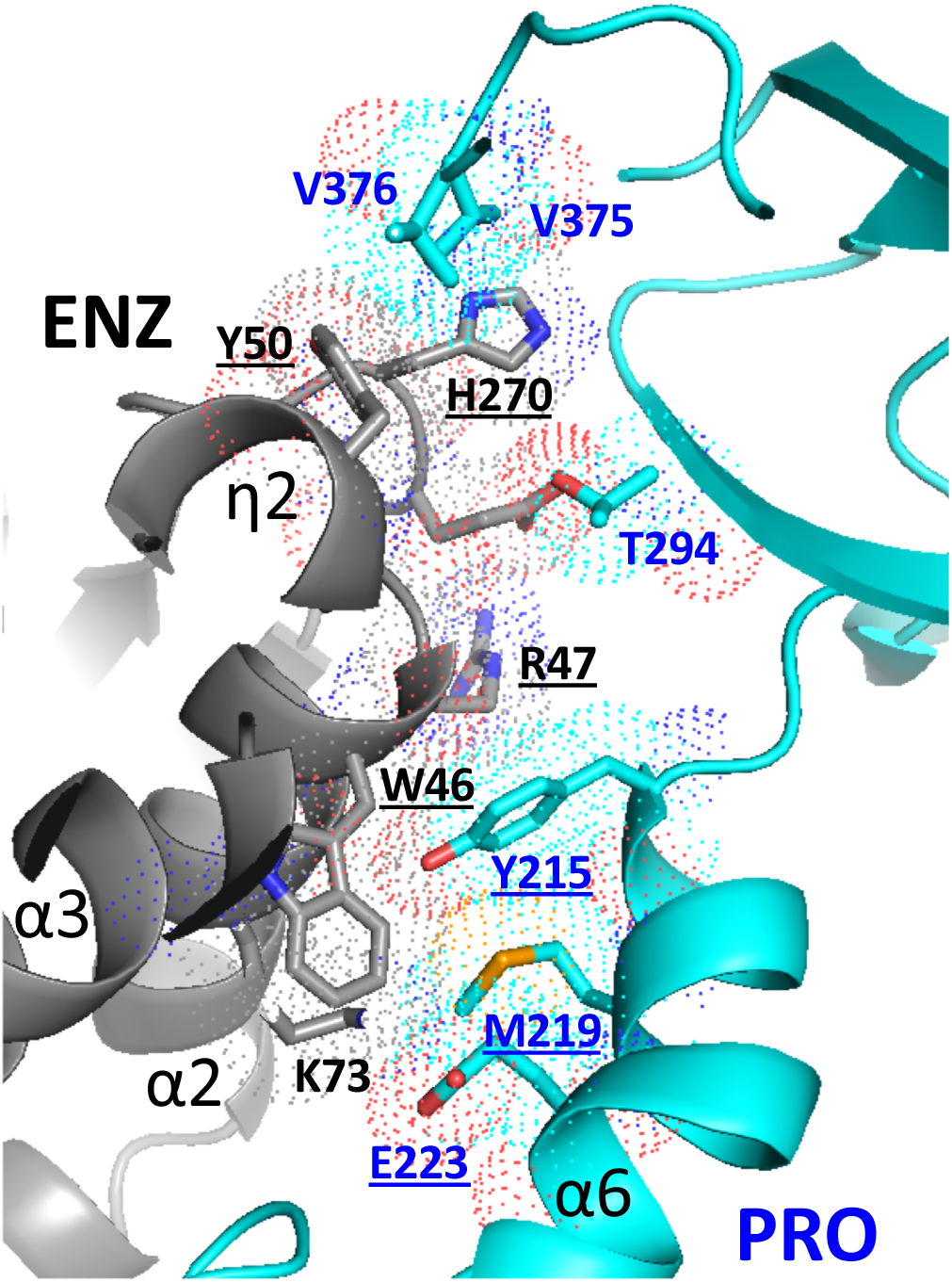
The interface between two ENZ-PRO heterodimers is dominated by van der Waals’ contacts. Residues forming key contacts between two ENZ-PRO heterodimers are displayed in dotted-sphere representation, indicating their van der Waals’ radii. Mutation of the underlined residues breaks the heterotetrameric assembly into ENZ-PRO heterodimers.

**Figure 6.**
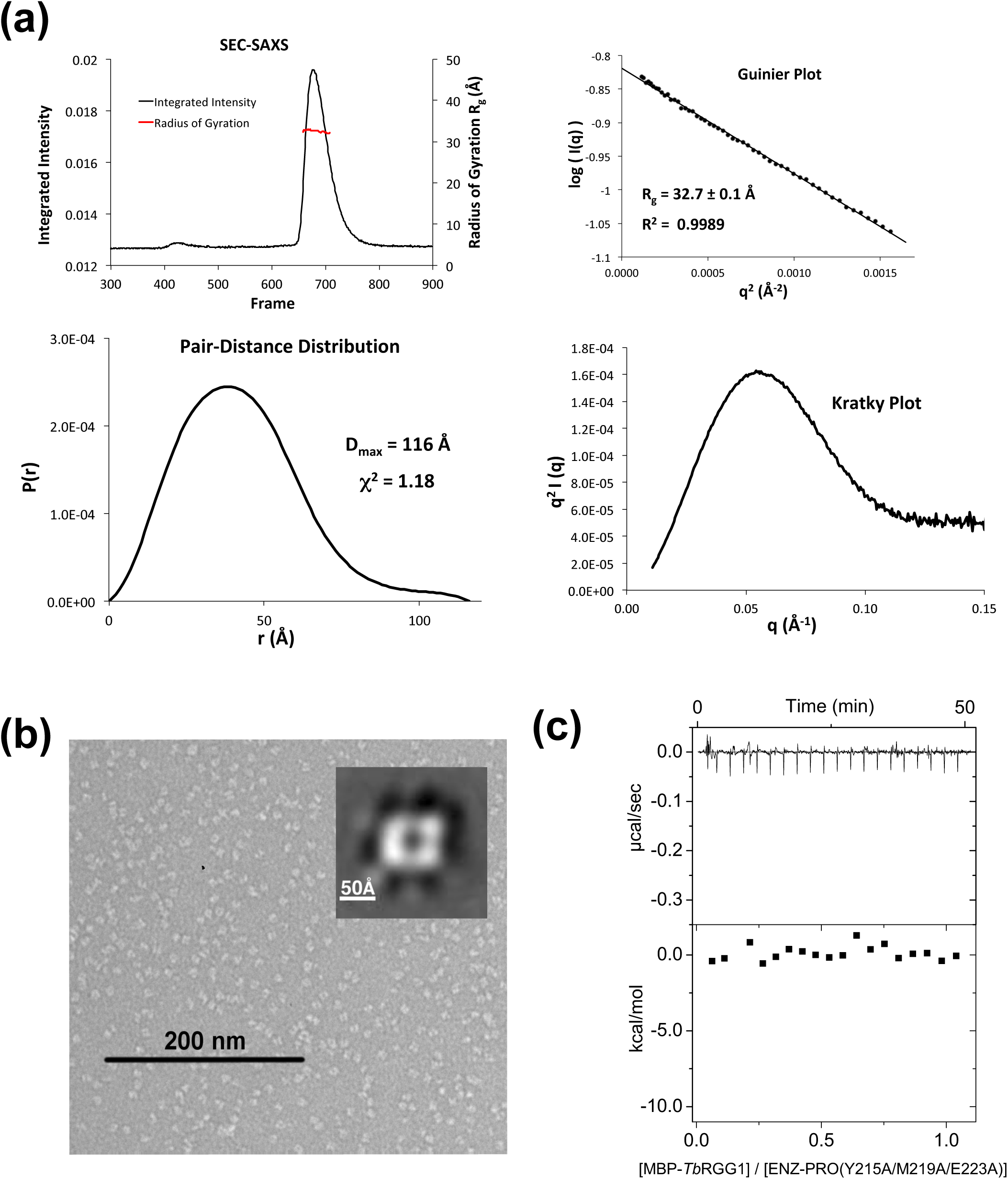
Structure-based mutagenesis of the tetrameric interface yields stable ENZ-PRO heterodimers. (a) SEC–SAXS analysis of a representative heterodimeric *Tb*PRMT1 ENZ-PRO mutant (the ENZ W46A/R47A/Y50A triple mutant). Top left panel: SEC-SAXS integrated intensities (left y-axis) plotted against frame number (x-axis). The red dots indicate radius of gyration, R_g_ (on the right y-axis). Top right panel: Guinier plot calculated from averaging buffer-subtracted scattering intensities. The coefficient of determination, R^2^, is 0.9989. Bottom left panel: Pair-distance distribution function P(r), yielding a maximum molecular diameter of 116 Å. Bottom right panel: Normalized Kratky plot calculated from SEC–SAXS data. (b) Negative-stain electron microscopy. EM micrograph with a 200 nm scale bar. Inset: Predominant 2D class average. (c) ITC thermogram (upper panel) and plots of corrected heat values (lower panel) for binding of the ENZ W46A/R47A/Y50A triple mutant to Maltose Binding Domain (MBP)-fused *Tb*RGG1 protein.

Moreover, the combination of a single mutation in each protein, H270A in ENZ and Y215A in PRO, disrupted the tetramer as well (**Tables 2** and **3**). In contrast to the rhomboid shape of the wild-type, negative-stain EM analysis of two mutants revealed a square-shaped structure, consistent in shape and size with an ENZ-PRO heterodimeric unit (**Figs. 3b** and **6b**). Extensive SEC-SAXS data obtained from these mutants confirm the substantially smaller size of the heterodimer with respect to the heterotetrameric wild-type structure (**Fig. 6a, Table 2**). These solution studies verify the dimer-dimer interface observed in the crystal structure. Importantly, ITC binding and methyltransferase assays showed that none of the dimeric mutants bound MBP-*Tb*RGG1 and did not possess any detectable methyltransferase activity (**Fig. 6c** and **Table 3**). On the other hand, the single mutations K73A, W64A, R47A, Y50A, and H270A in ENZ, the double alanine mutant H270A/E271A in ENZ, single mutations Y215A and K232A in PRO retained the tetrameric assembly and methylate the substrate as the wild-type complex (**Table 3**). We conclude that the determined ENZ-PRO heterotetramer structure is the biological functional unit both *in vitro* and *in vivo* [14], and that heterotetramer assembly is required for substrate binding and methyltransferase activity.

## Discussion

Catalytically inert pseudoenzymes are abundant in the proteome and perform diverse functions, serving as scaffold proteins [40], modulators of enzyme activity and signaling pathway components [26], or competitors of active paralogs for substrate(s) [41]. Prozymes are a subgroup of pseudoenzymes that specifically stimulate an otherwise inactive enzyme by complex formation. Here, we describe a chaperone function of the *Tb*PRMT1 prozyme (PRO) in complex with the *Tb*PRMT1 enzyme (ENZ) in *T. brucei*. Structural and functional analyses reveal distinct structural features that set PRO apart from its active counterpart ENZ and elucidate the mechanism of *Tb*PRMT1 ENZ activation by PRO through oligomerization.

Sequence analysis alone had already indicated that PRO was lacking conserved residues that are critical for catalysis and methyl donor (AdoMet) binding and hence provided evidence that PRO would catalytically be inactive [14]. Indeed, the crystal structure of the ENZ-PRO complex showed that the cofactor product was only bound to ENZ, in agreement with an AdoMet crosslink experiment (**Fig. 3h**) [14]. However, our structural analysis revealed further marked differences on the secondary-structure level of PRO with respect to ENZ (**Fig. 1**), most notably the lack of the η2 3_10_-helix in the Rossmann fold of PRO, which affect its tertiary structure (**Fig. 3g**) and which are ultimately critical for its catalytic inactivity. The lack of the η2 3_10_-helix results in a tilt of several adjacent *α*-helices, which in turn affect dimerization and abolish cofactor binding in the prozyme (**Fig. 3g**).

While ENZ possesses all essential residues and secondary structure elements for cofactor AdoMet binding and catalysis (**Figs. 1** and **3h**), ENZ itself is unstable in solution in the absence of PRO and hence incapable of catalysis [23]. ENZ features highly hydrophobic patches in the dimerization arm that are thermodynamically unfavorable if exposed to solvent [42]. While hydrophobic patches are also found in other PRMTs, homodimerization covers these hydrophobic patches to stabilize the active enzymes [8, 17–19]. By contrast, ENZ homodimerization is sterically prevented, specifically by Tyr190 within the dimerization arm of ENZ (**Fig. 4e**). In mammalian and yeast class I PRMTs, as well as in *Tb*PRMT1 PRO, the corresponding residue is mostly a cysteine or a proline, thus facilitating formation of a stable homodimer [14, 17–19] (**Fig. 4a**). However, we note that the Tyr190 mutation to Cys or Pro was not sufficient to form homodimeric enzyme complex (data not shown), suggesting that homodimerization does not depend on a single residue.

Higher-order oligomerization beyond the homodimer is well established in PRMTs. For example, PRMT1s from various organisms form a hexamer in solution [17, 19–21]. While the oligomerization of mammalian and yeast PRMT1s is dependent on PRMT concentration [17], we did not detect such a dependence for the *Tb*PRMT1 complex, as it always remained tetrameric under different protein concentrations (**Supplemental Fig. S5**). Dimerization is required for AdoMet binding in PRMTs [17–19], and even higher oligomerization of PRMT1 is required for its catalysis *in vivo* [39]. The dimeric fractions of human PRMT1 did not show methyltransferase activity, while the higher oligomeric fractions did. As we demonstrated here, substrate binding and methyltransferase activities are essentially abolished in the stable *Tb*PRMT1 ENZ-PRO dimeric complex mutants with respect to the wild-type tetrameric complex (**Table 2**). We conclude that oligomers beyond the dimer may generally constitute the active species of PRMT1s, with dimerization not being sufficient for its activities *in vitro* and/or *in vivo*.

Although the ENZ-PRO heterotetramer possesses two active sites, only one MBP-*Tb*RGG1 substrate molecule is bound to the tetramer (**Fig. 2c**). In principle, an allosteric mechanism between the two heterodimers could explain this finding, whereby substrate binding to one heterodimer would induce changes in the other dimer that would prevent further substrate binding. Alternatively, a tetramer could provide a unique composite interaction surface not present in the heterodimer. Our ITC experiments show that substrate binding is abolished in heterodimer mutants (**Table 2**), supporting the latter mechanism. We cannot exclude though that both ENZ-PRO heterodimers of a tetramer may independently and in parallel be engaged in the methylation of other substrate proteins or even *Tb*RGG1 without the bulky MBP fusion partner [13]. However, attempts to remove the MBP fusion partner from *Tb*RGG1 have not yielded stable *Tb*RGG1.

The concept that PRO is necessary to form a stable, catalytically active PRMT heterodimer with ENZ has important implications for regulation. In essence, the activity of *Trypanosoma* PRMT1 would be regulated by the protein amount of *Tb*PRMT1 prozyme. As the knock down of *Tb*PRMT1 prozyme has shown, the its mRNA reduction did not affect the mRNA amount of *Tb*PRMT1 ENZ, but affected the protein amount of *Tb*PRMT1 ENZ [23]. Unlike all previously reported PRMTs, *Tb*PRMT1 ENZ is unstable on its own because it cannot form a homodimer, in part due to Tyr190 of *Tb*PRMT1 ENZ by steric clashes (**Fig. 4e**). PRO, but not ENZ, was found in stress granules, where it would not be accessible to ENZ translated in the cytosol, implying that sequestration of PRO may provide a means of controlling ENZ activity. We propose that PRO serves as a folding chaperone for its catalytic partner, providing a new paradigm for prozyme function. Thus, the expression and localization of PRO are ultimately determinants of *Tb*PRMT1 ENZ activity, Sequence alignments of ENZ and PRO with *Tb*PRMT1 of related protozoan kinetoplastids (**Supplemental Figs. S6 and S7**) including the human parasites *T. cruzi* causing Chagas disease and *Leishmania donovani* causing visceral leishmaniasis (black fever) suggest that ENZ-PRO complexes also exist in many other parasites and corroborate numerous features and conclusions that we present here for *Tb*PRMT1. Among the putative PRO homologs, AdoMet binding residues are not conserved, and the residues forming the η2 3_10_-helix are missing, consistent with catalytic inactivity of PRO. Furthermore, dimerization residues of ENZ and PRO are vastly conserved, indicating the same heterodimer formation in other kinetoplastids. Specifically, Tyr190 of ENZ is invariant, arguing that the ENZ proteins of other species are similarly incapable of homodimerization because of steric clashes and that they therefore necessitate a prozyme for stability. Intriguingly, tetrameric interface residues are largely conserved as well, which provides further evidence that heterotetramers are the active species and a prerequisite for methylation. Finally, the first 40 N-terminal PRO residues are not conserved, which coincides with our finding that this region is dispensable for substrate binding, while the adjacent residues 41-52 are conserved and critical for substrate recognition. Among the kinetoplastid PRMTs, the enzyme-prozyme paradigm only exists for PRMT1, while *Tb*PRMT5, *Tb*PRMT6 [43], and *Tb*PRMT7 [9, 10] do not have such prozymes for regulation.

From an evolutionary standpoint, we speculate that ENZ and PRO have co-evolved to furnish a functional PRMT1 enzyme, as a mutation in ENZ such as Cys190-to-Tyr would render ENZ unstable, unless PRO concomitantly emerged to function as a folding chaperone for ENZ. Initially, PRO may have been catalytically active, but over time, mutations of the PRO AdoMet binding site may have transformed PRO into a catalytically dead enzyme, focusing on its primary role as a regulator of ENZ. Even if the AdoMet binding residues were not mutated, lack of η2 3_10_-helix alone may have compromised AdoMet binding within the Rossmann fold. The fact that PRMT1 ENZ-PRO complexes are conserved throughout kinetoplastids suggests that this regulatory mechanism proved to be valuable to these organisms and may constitute a general mechanism of PRMT regulation beyond kinetoplastids.

## Materials and Methods

### Protein expression and purification

DNA fragments of *Tb*PRMT1^PRO^ (TriTrypDB: **<u>Tb927.10.3560</u>**) and *Tb*PRMT1^ENZ^ (TriTrypDB: **<u>Tb927.1.4690</u>**) were amplified by PCR from genomic DNA and cloned into the multiple coning sites (MCS) 1 and 2 using the NcoI/NotI and NdeI/XhoI restriction sites of a modified pETDuet-1 vector (Novagen) containing an N-terminal PreScission protease (GE Healthcare) cleavable His_6_-tag prior to MCS 1. The constructs were overexpressed in *E. coli* BL21-CodonPlus(DE3)-RIL cells (Stratagene) and grown in LB media containing appropriate antibiotics. Mutations in ENZ and PRO were introduced by overlap extension PCR mutagenesis. Protein expression was induced at OD_600_ of ≈0.4 with 0.1 mM isopropyl-*β*-D-thiogalactoside at 18 °C for 16 h. The cells were harvested by centrifugation at 7,500 x *g* and 4°C and lysed with a cell disrupter (Avestin) in a buffer containing 20 mM Tris, pH 8.0, 300 mM NaCl, 5 mM *β*-mercaptoethanol, 0.5 mM 4-(2-aminoethyl)benzenesulfonyl fluoride hydrochloride (Sigma), 2 μM bovine lung aprotinin (Sigma), and complete EDTA-free protease inhibitor cocktail (Roche). After centrifugation at 35,000 x *g* for 45 min, the cleared lysate was loaded onto a Ni-NTA column (Qiagen) and eluted with an imidazole gradient. Protein-containing fractions were pooled, dialyzed against a buffer containing 20 mM Tris, pH 8.0, 100 mM NaCl, and 5 mM dithiothreitol (DTT), and subjected to cleavage with PreScission protease (GE Healthcare) for 5 h at 4°C. Following His_6_-tag removal, the cleaved protein was bound to a heparin column (GE Healthcare) and eluted with a NaCl gradient. Protein-containing fractions were pooled, concentrated, and purified on a HiLoad Superdex 200 16/60 gel filtration column (GE Healthcare) in a buffer containing 20 mM HEPES, pH 7.5, 150 mM NaCl, and 0.5 mM Tris(2-carboxyethyl)phosphine hydrochloride (TCEP).

MBP-tagged *Tb*RGG1 was cloned into a pMAL-c2X vector (NEB) using BamHI and SalI restriction sites. *Tb*RGG1 was overexpressed in *E. coli* BL21-CodonPlus(DE3)-RIL cells (Stratagene) and grown in LB media containing appropriate antibiotics. Protein expression was induced at OD_600_ of ≈0.4 with 0.1 mM isopropyl-*β*-D-thiogalactoside at 18 °C for 16 h. The cells were harvested by centrifugation at 7,500 x *g* and 4°C and lysed with a cell disrupter (Avestin) in a buffer containing 20 mM Tris, pH 8.0, 300 mM NaCl, 5 mM *β*-mercaptoethanol, 0.5 mM 4-(2-aminoethyl)benzenesulfonyl fluoride hydrochloride (Sigma), 2 μM bovine lung aprotinin (Sigma), and complete EDTA-free protease inhibitor cocktail (Roche). After centrifugation at 35,000 x *g* for 45 min, the cleared lysate was loaded onto a amylose resin (NEB) and eluted with a maltose gradient. Protein-containing fractions were pooled, dialyzed against a buffer containing 20 mM Tris, pH 7.5, 20 mM NaCl, and 5 mM dithiothreitol (DTT). The protein was bound to a SP column (GE Healthcare) and eluted with a NaCl gradient. Protein-containing fractions were pooled, concentrated, and purified on a HiLoad Superdex 200 16/60 gel filtration column (GE Healthcare) in a buffer containing 20 mM HEPES, pH 7.5, 150 mM NaCl, and 0.5 mM TCEP.

### Crystallization, data collection, structure determination and refinement

For formation of the complex with the AdoHcy, 4.5 mg/ml of purified ENZ-PROΔ52 was mixed in a 1:8 molar ratio with AdoHcy and incubated for 4 h on ice. The crystallization solution consisted of 7% PEG 4000 and 0.1 M Tris, pH 7.4. Crystals grew in space group C2 at room temperature within a week. X-ray diffraction data were collected at the 24ID-C beamline at the NE-CAT at the Advanced Photon Source (APS) of Argonne National Laboratory (ANL). Diffraction data were processed in HKL2000 [44]. The structure was solved by the single anomalous dispersion (SAD) phasing technique in the program AutoSol of the PHENIX package [45], using data obtained from seleno-L-methionine-labeled crystals. The asymmetric unit contained one tetramer. Model building was performed in O [46] and Coot [47]. The final model spanning residues 38-384 was refined in Phenix [45] to an R_free_ of 26.1 % with excellent stereochemistry as assessed by MolProbity [48]. Details for data collection and refinement statistics are summarized in **Supplemental Table S1**. Figures were generated using PyMOL (Schrödinger, LLC), the electrostatic potential was calculated with APBS [49]. Atomic coordinates and structure factors have been deposited with the Protein Data Bank under PDB: **<u>6DNZ</u>**.

### Multi-angle light scattering

Purified proteins were characterized by multi-angle light scattering following size-exclusion chromatography [50]. Protein at 50 µM was injected onto a Superdex 200 10/300 GL size-exclusion chromatography column (GE Healthcare) equilibrated in a buffer containing 20 mM HEPES pH 7.5, 150 mM NaCl, and 0.5 mM TCEP. The chromatography system was connected in series with an 18-angle light scattering detector (DAWN HELEOS) and refractive index detector (OptilabrEX) (Wyatt Technology). Data were collected every second at a flow rate of 0.25 mL/min at 25 °C. Data analysis was carried out using the program ASTRA, yielding the molar mass and mass distribution (polydispersity) of the sample.

### Isothermal titration calorimetry

ITC measurements were performed at 25°C using a MicroCal auto-iTC200 calorimeter (MicroCal, LLC). Wild-type and mutant ENZ-PRO proteins as well as MBP-*Tb*RGG1 protein were extensively dialyzed against a buffer containing 20 mM HEPES, pH 7.5, 150 mM NaCl, and 0.5 mM TCEP. 2 µL of 0.13 mM MBP-*Tb*RGG1 was injected into 0.2 mL of 0.03 mM ENZ-PRO proteins in the chamber every 150 s. Baseline-corrected data were analyzed with ORIGIN software.

### Methylation assays

To assay the activity of *Tb*PRMT1 tetramerization mutants, 37.5 nM *Tb*PRMT1 tetramer was mixed with 6 μM MBP-TBRGG substrate, 0.7 μM [^3^H]AdoMet (Adenosyl-L-Methionine, S-[methyl-^3^H]-, 55-85Ci (2.03-3.15TBq)/mmol; PerkinElmer), 9.3 μM unlabeled AdoMet, 2 mM DTT and 2mM PMSF in PBS in total volume of 25 μl. Reactions were incubated at 26°C for 1.5 hour, stopped by addition of SDS loading dye and separated on SDS PAGE. Gel was Coomassie stained and soaked in EN^3^HANCE (PerkinElmer). Dried gel was then exposed to film for one week at −80°C. To assay the activity of *Tb*PRMT1 containing N-terminal prozyme truncations, reactions were performed as above, except the amount of MBP-TBRGG substrate was lowered to 0.6 μM, unlabeled AdoMet was left out and 6 μg of MBP2* protein (NEB) was added to each reaction to increase molecular crowding.

### Small-Angle X-ray Scattering

SEC–SAXS of wild-type and mutant ENZ-PRO proteins was performed at the G1 station at MacCHESS, which is equipped with an ÄKTA Pure FPLC system (GE Healthcare). Protein was loaded at concentrations ranging from 2 to 16 mg/ml on a Superdex 200 10/300 GL column (GE Healthcare) equilibrated in 20 mM HEPES, pH 7.5, 150 mM NaCl, and 0.5 mM TCEP. SAXS data were recorded on a Pilatus 100 K-S detector at 2 s per frame with a fixed camera length of 1.522 m and an energy of 9.91 keV, allowing the collection of the angular range *q* of 0.01–0.30 Å^−1^. Primary reduction of the SAXS data was performed using RAW [51]. A Guinier plot of the buffer-subtracted profile was linear to the lowest measured *q* value. GNOM [52] was used to calculate *P*(*r*) plots from the scattering data. The maximum diameter was chosen so that the *P*(*r*) function fell gradually to zero at *r* = *D*_max_ unconstrained. Theoretical radii of gyration were calculated using CRYSOL [53]. SEC–SAXS data collection and analysis statistics are listed in **Table 2**.

### Electron Microscopy

Negative stain transmission electron microscopy was performed on the wild-type ENZ-PRO complex and the Y215A/M219A/E223A triple mutant. Samples at 0.03 mg/ml were stained using 2% uranyl formate on continuous carbon grids. Micrographs were collected on the JEOL-1230 transmission electron microscope with a Gatan US400 detector. Data were processed using the Appion pipline and CryoSPARC [54, 55]. Using Appion, CTF estimation was performed using CTFFIND4 [56]. Automated particle picking was done using DoG Picker [57]. An initial stack of particles was assembled in Appion. 2D Classification and 3D refinement were performed in CryoSPARC.

## Accession numbers

The x-ray structure (coordinates and structure files) of the *Tb*PRMT1 ENZ-Δ52PRO complex with AdoHcy have been deposited in the PDB with accession number **<u>6DNZ</u>**.

## Acknowledgements

We thank David King (University of California, Berkeley) for mass spectrometry analysis, Igor Kourinov and Kay Perry for support during data collection at NE-CAT, Richard Gillilan for support during data collection at CHESS, the High-Throughput Screening and Spectroscopy Resource Center at Rockefeller University, and the X-Ray Crystallography & Molecular Interactions at the Sidney Kimmel Cancer Center, which is supported in part by National Cancer Institute Cancer Center Support Grant P30 CA56036 and S10 OD017987. X-ray data were collected at the NE-CAT beamlines (GM103403) on a Pilatus detector (RR029205) at the APS (DE-AC02-06CH11357). CHESS is supported by the NSF award DMR-1332208, and the MacCHESS resource is supported by NIGMS award GM-103485. The electron microscopy work was performed at the Simons Electron Microscopy Center and National Resource for Automated Molecular Microscopy located at the New York Structural Biology Center, supported by grants from the Simons Foundation (SF349247), NYSTAR, and the NIH National Institute of General Medical Sciences (GM103310). This work was supported by National Institutes of Health Grant R01 AI060260 (to L. K. R.) and American Heart Association Predoctoral Fellowship 15PRE24480155 (to L. K.). We would like to thank the members of the Debler laboratory for helpful discussions.

## Supplemental Information

**Figure S1.**
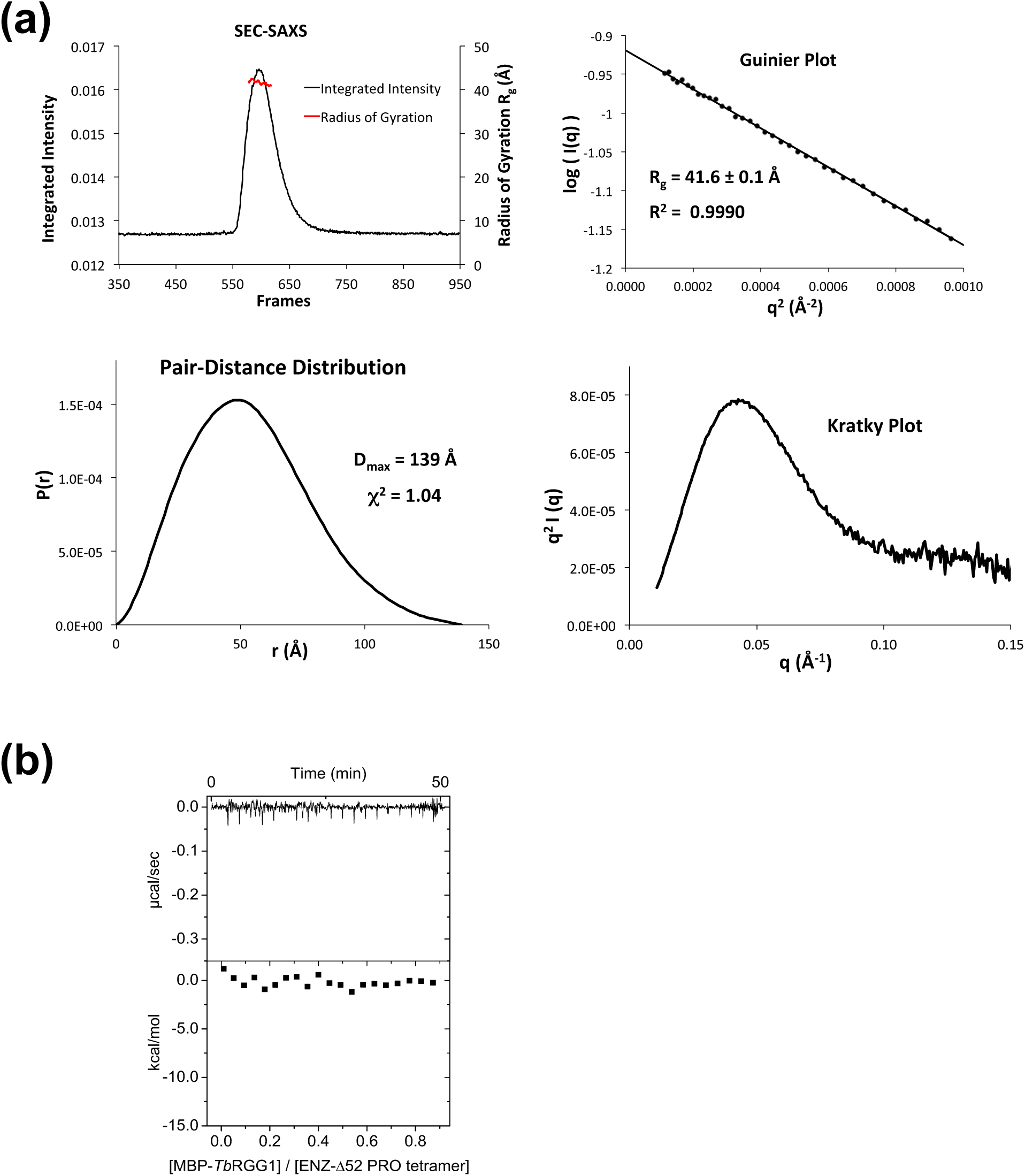
ENZ-Δ52PRO forms a tetramer in solution, but does not bind substrate. **(a)** Small-angle x-ray scattering analysis. Upper left: SEC-SAXS profile (integrated intensity vs. frames). The red dots indicate R_g_ values (on the right y-axis) corresponding to frames. Upper right: Guinier plot. Lower left: pair-distance distribution, and Kratky plot (lower right). **(b)** ITC analysis of ENZ-Δ52PRO titrated with MBP-*Tb*RGG1.

**Figure S2.**
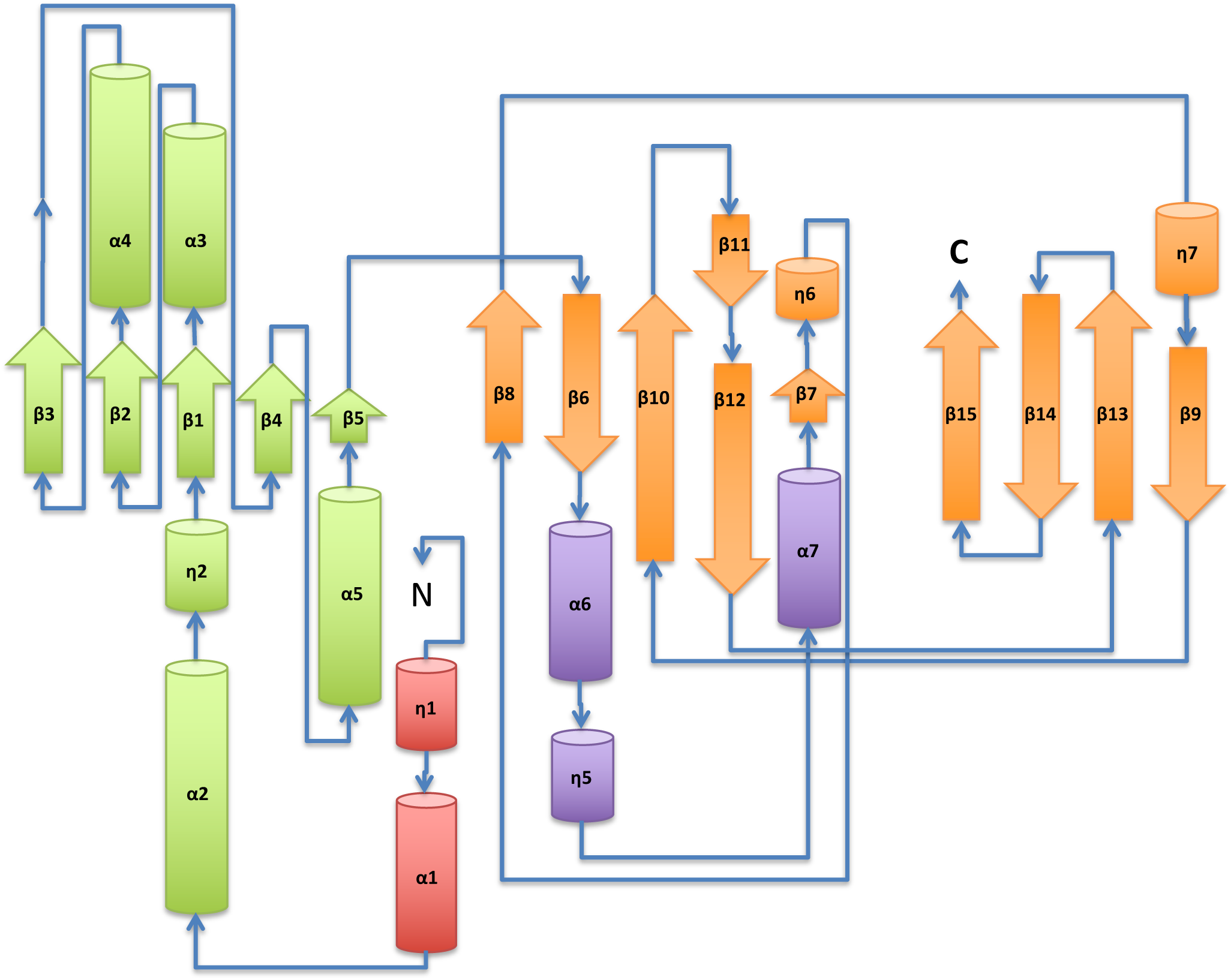
Topology of *Rattus norvegicus* PRMT1 (PDB: 1OR8).

**Figure S3.**
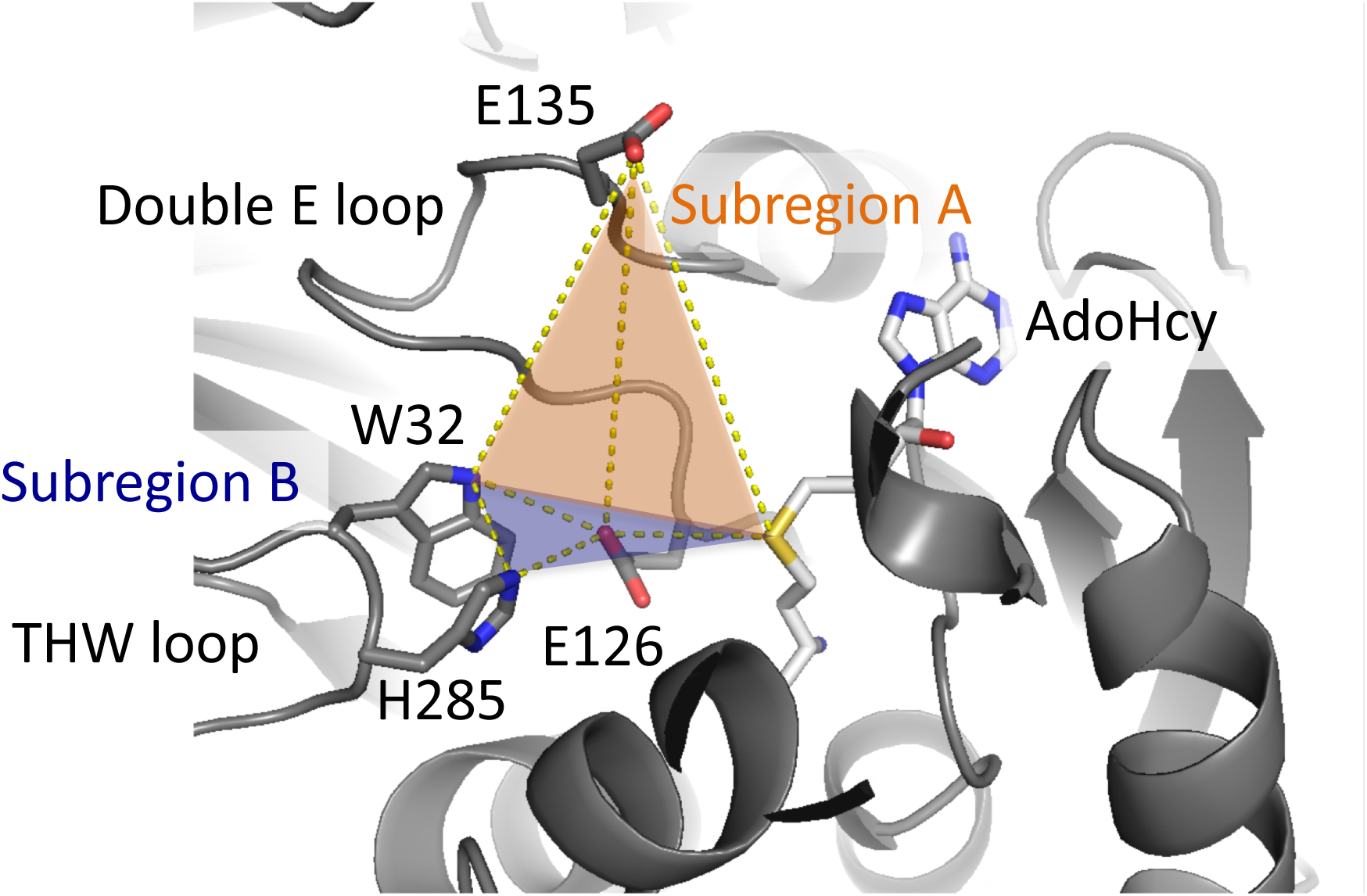
ENZ features a PRMT type I active-site architecture.

**Figure S4.**
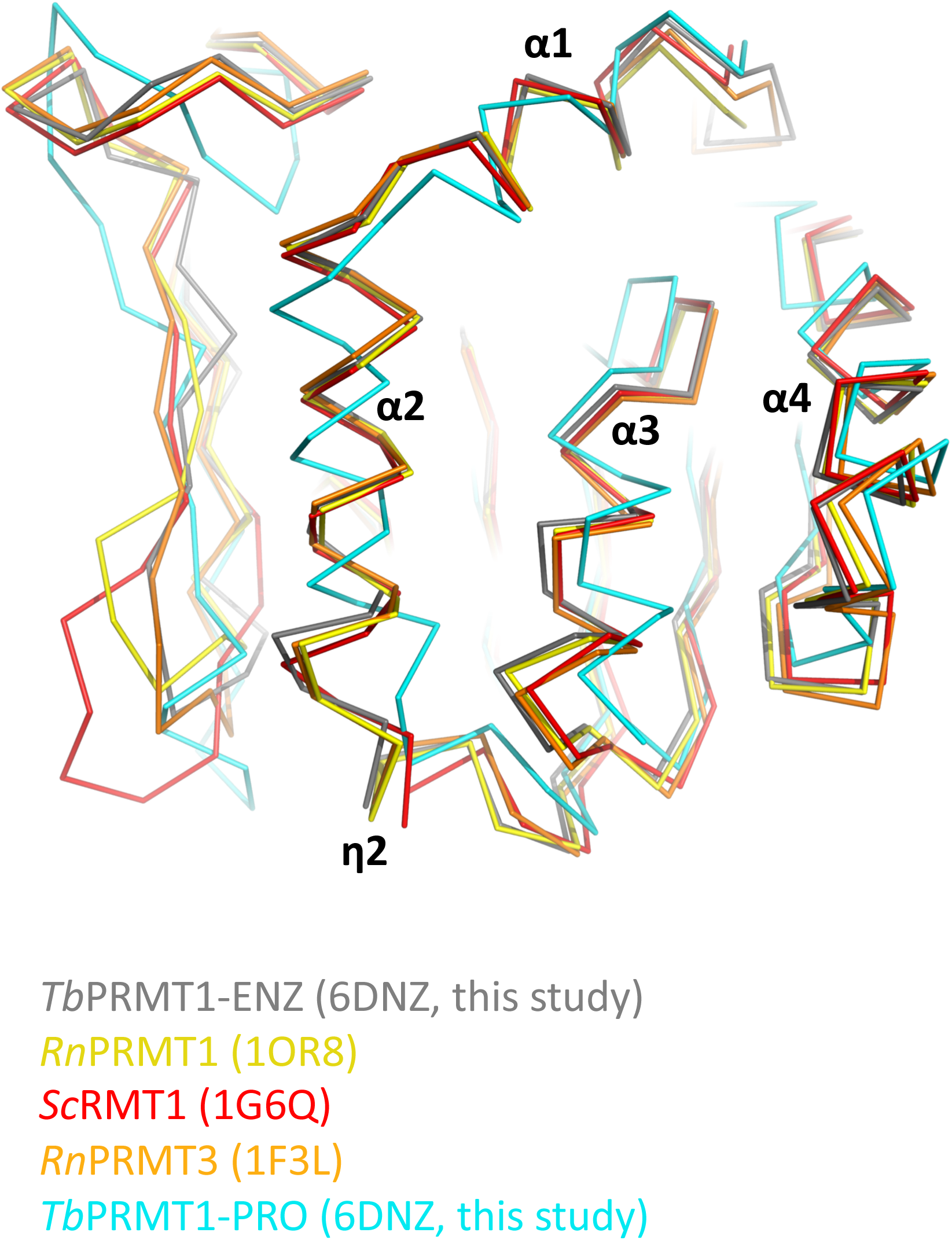
Superimposition of the Rossmann fold of various PRMT proteins. *Tb*: *Trypanosoma brucei*, *Rn*: *Rattus norvegicus*, *Sc*: *Saccharomyces cerevisiae*.

**Figure S5.**
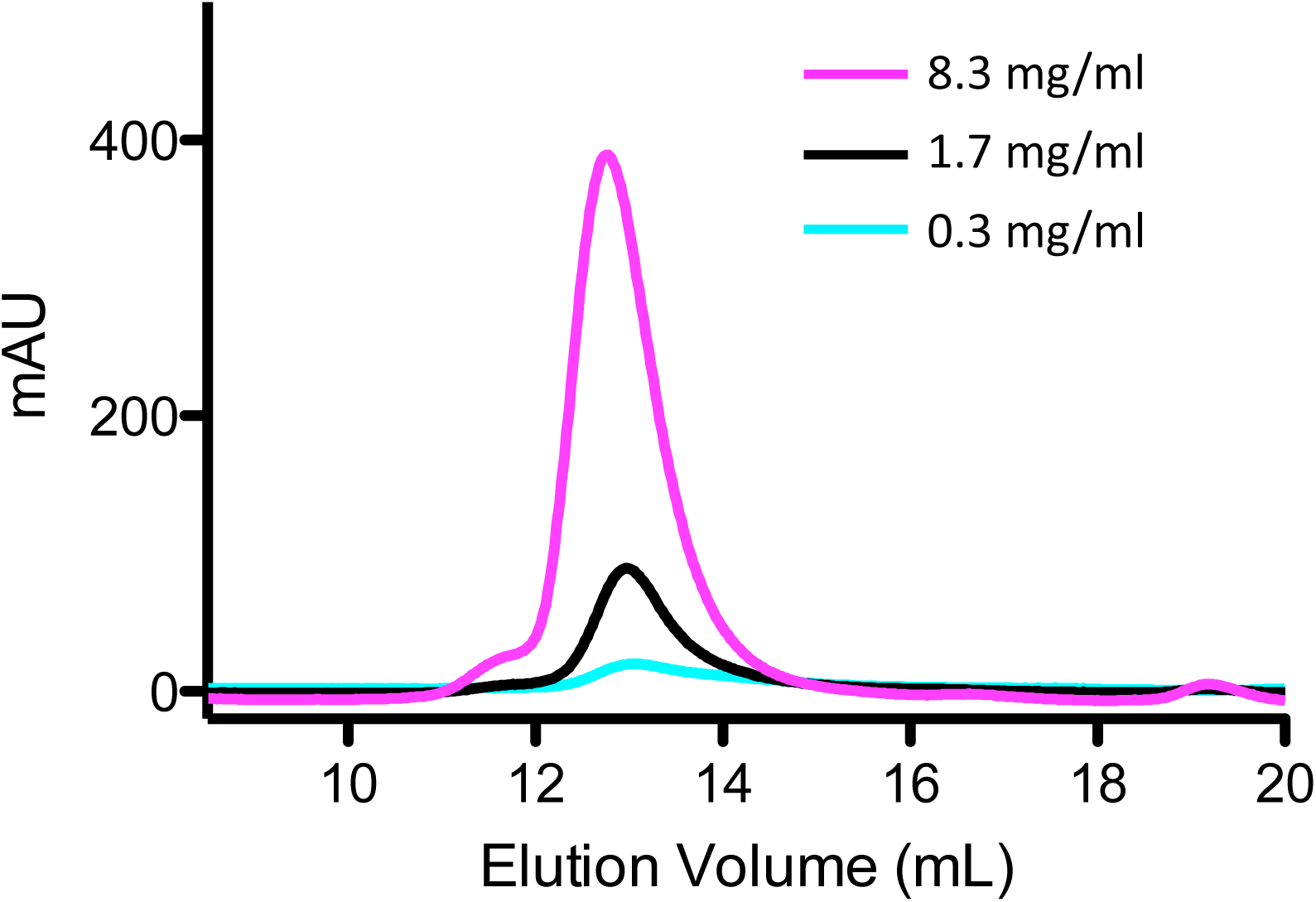
Concentration-independent heterotetramerization of ENZ-PRO. Size-exclusion chromatography (Superdex 200 10/300 column; GE Healthcare) profiles of ENZ-PRO at different concentrations. Absorbance was recorded at 280 nm.

**Figure S6.**
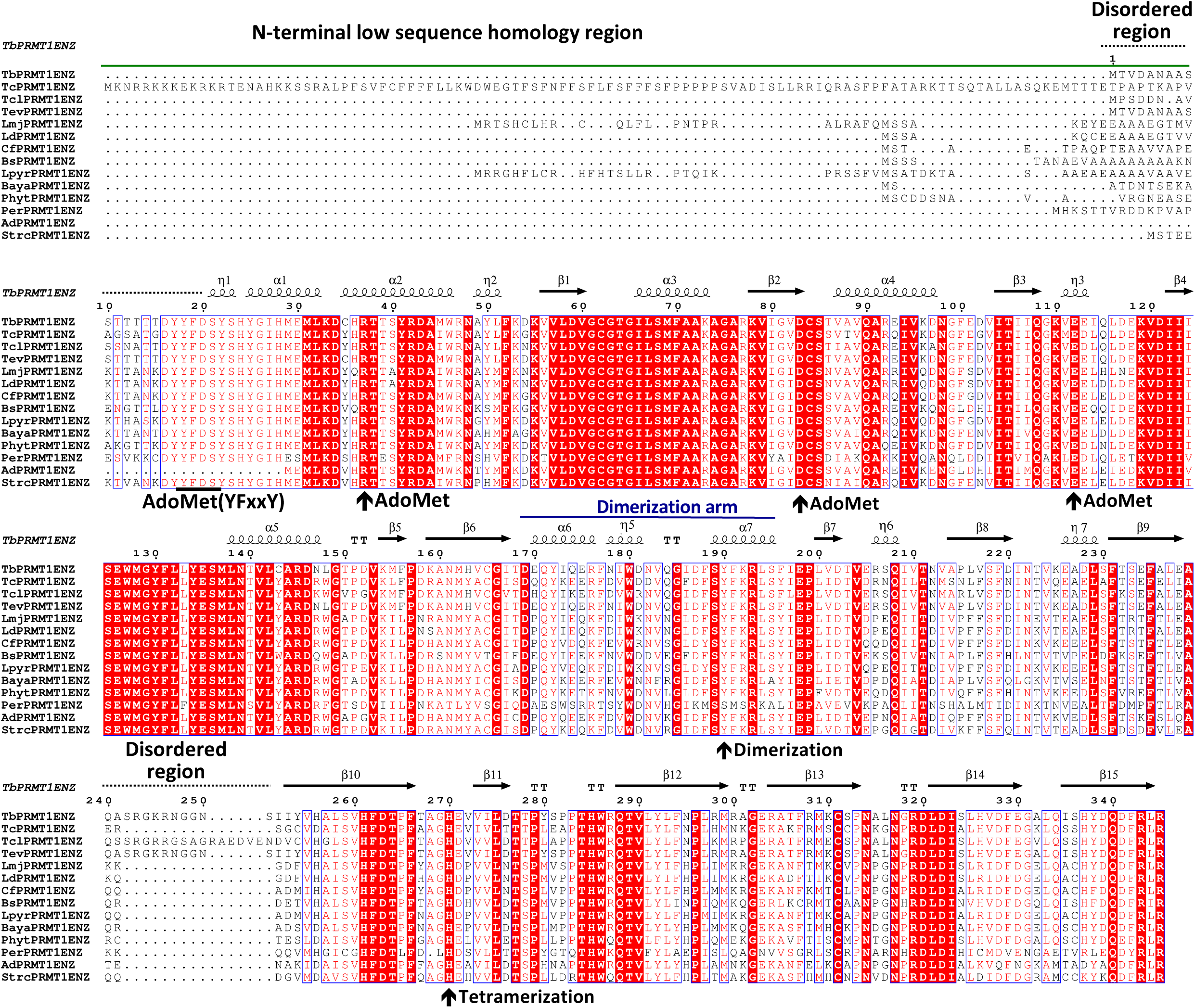
Multiple sequence alignment of putative PRMT1 enzymes among kinetoplastids. The N-terminal low sequence homology region, secondary structural elements, the dimerization arm, and disordered regions are indicated above the alignment. Key residues for AdoMet binding, dimerization, and tetramerization of *Tb*PRMT1 ENZ are indicated by arrows below the sequence. Tb: *Trypanosoma brucei* (TriTrypDB: Tb927.1.4690), Tc: *Trypanosoma cruzi* (TriTrypDB ID: TcCLB.506529.50), Tcl: *Trypanosoma congolense* (TriTrypDB ID: TcIL3000_1_1890), Tev: *Trypanosoma evansi* (TriTrypDB ID: TevSTIB805.1.4590), *Trypanosoma vivax* (TriTrypDB ID: TvY486_1003550), Ld: *Leishmania donovani* (TriTrypDB ID: LdBPK_120850.1), Cf: *Crithidia fasciculata* (TriTrypDB ID: CFAC1_010018100), Bs: *Bodo saltans* (GenBank ID: CUG92929.1), Lpyr: *Leptomonas pyrrhocoris* (TriTrypDB ID: LpyrH10_23_0130), Baya: *Blechomonas ayalai* (TriTrypDB ID: Baya_031_0090), Per: *Perkinsela sp.* (GenBank ID: KNH05275.1), Ad: *Angomonas deanei* (GenBank ID: EPY25076.1), and Strc: *Strigomonas culicis* (GenBank ID: EPY24636.1).

**Figure S7.**
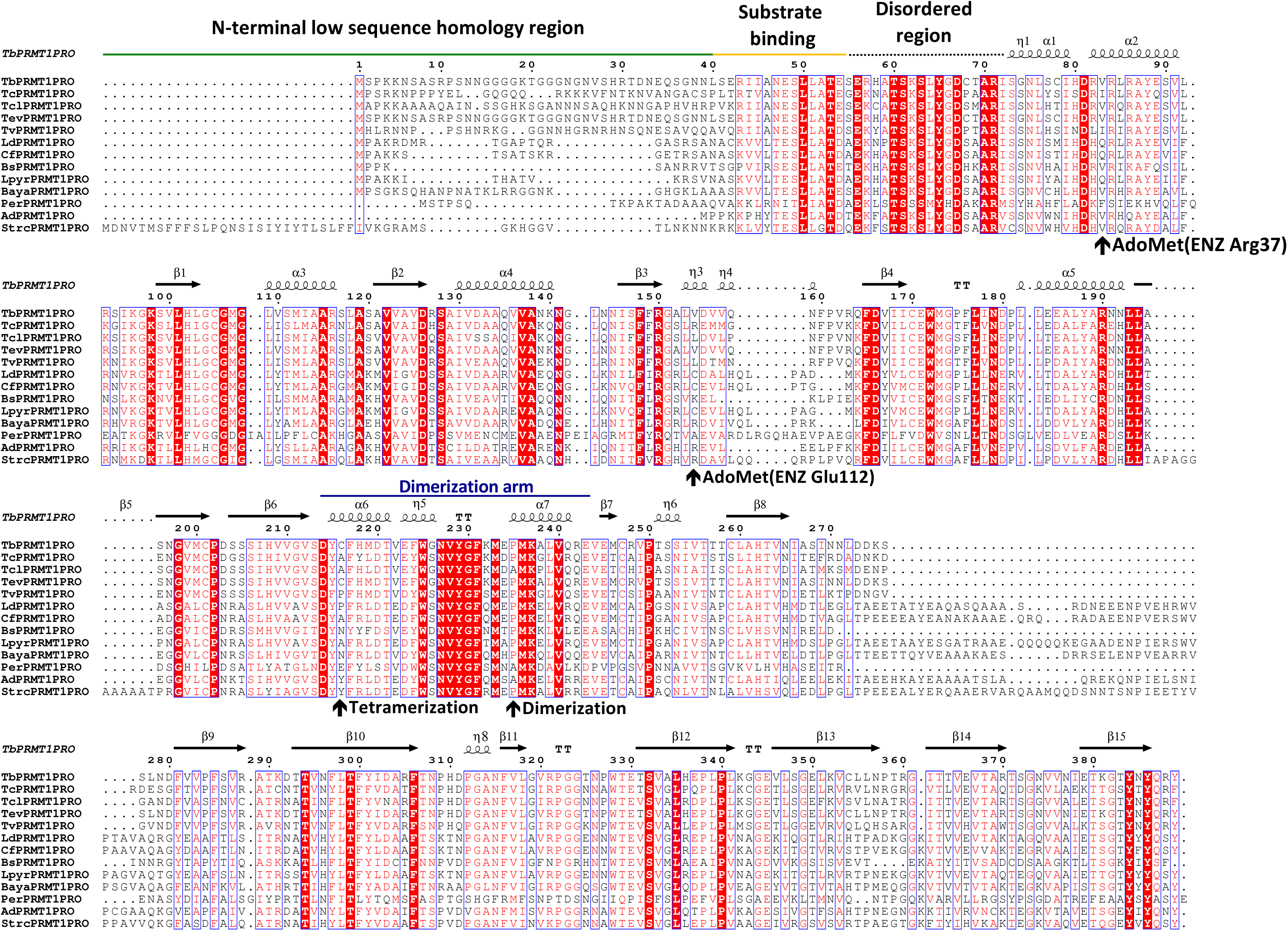
Sequence alignment of putative PRMT1 prozymes among kinetoplastids. Secondary structural elements, the N-terminal low sequence homology region, the substrate binding region, and the disordered region are indicated above the alignment. Key residues for dimerization, tetramerization, and lack of AdoMet binding of *Tb*PRMT1 PRO are indicated by arrows below the sequence. Tb: *Trypanosoma brucei* (TriTrypDB: Tb927.10.3560), Tc: *Trypanosoma cruzi* (TriTrypDB ID: TcCLB.510311.140), Tcl: *Trypanosoma congolense* (TriTrypDB ID: TcIL3000_10_2970), Tev: *Trypanosoma evansi* (TriTrypDB ID: TevSTIB805.10.3790), *Trypanosoma vivax* (TriTrypDB ID: TvY486_1003550), Ld: *Leishmania donovani* (TriTrypDB ID: LdBPK_030580.1), Cf: *Crithidia fasciculata* (TriTrypDB ID: CFAC1_060012600), Bs: *Bodo saltans* (GenBank ID: CUG88155.1), Lpyr: *Leptomonas pyrrhocoris* (TriTrypDB ID: LpyrH10_34_0780), Baya: *Blechomonas ayalai* (TriTrypDB ID: Baya_055_0080), Per: *Perkinsela sp.* (GenBank ID: KNH01754.1), Ad: *Angomonas deanei* (GenBank ID: EPY40540.1), and Strc: *Strigomonas culicis* (GenBank ID: EPY28765.1).

**Supplemental Table S1.**
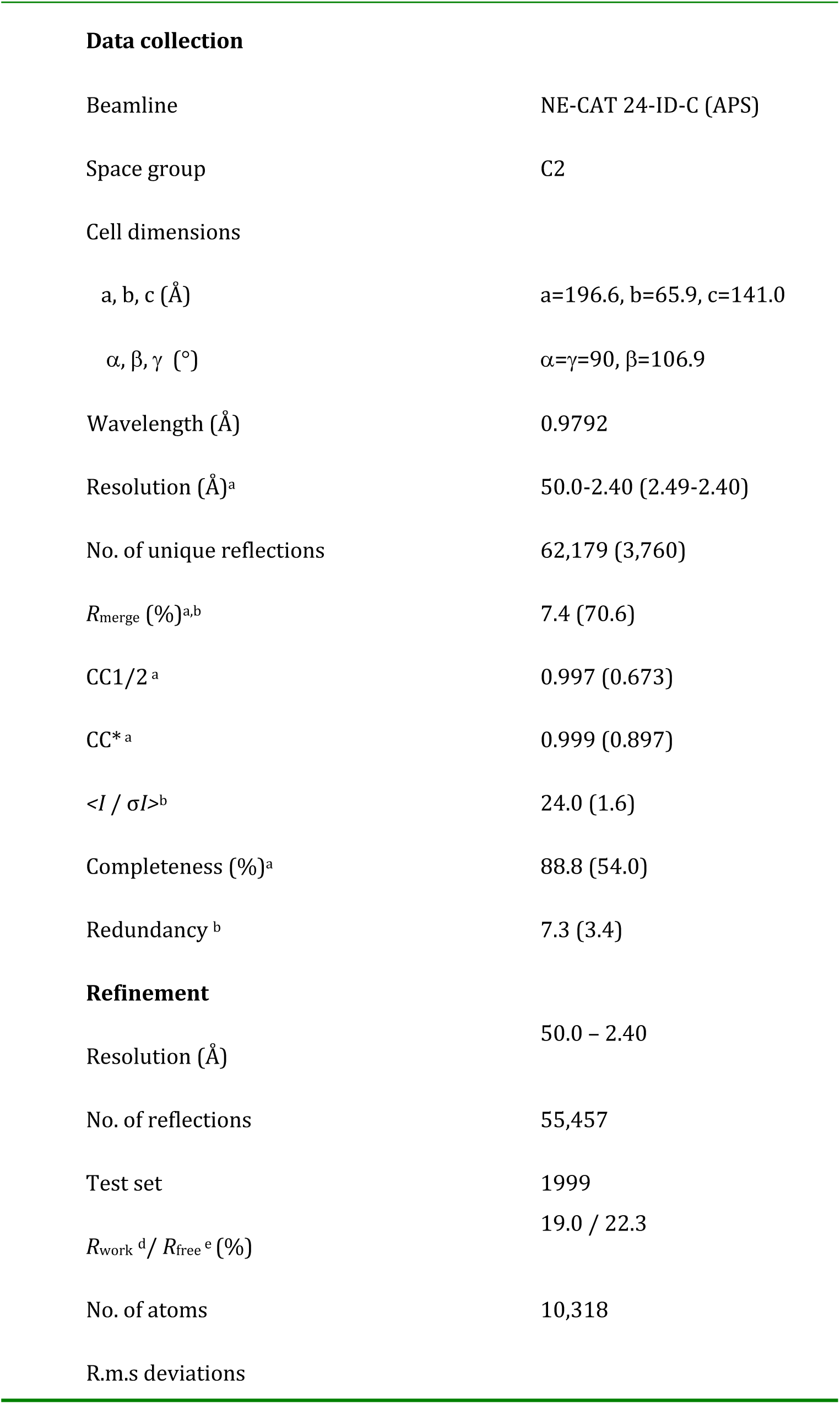
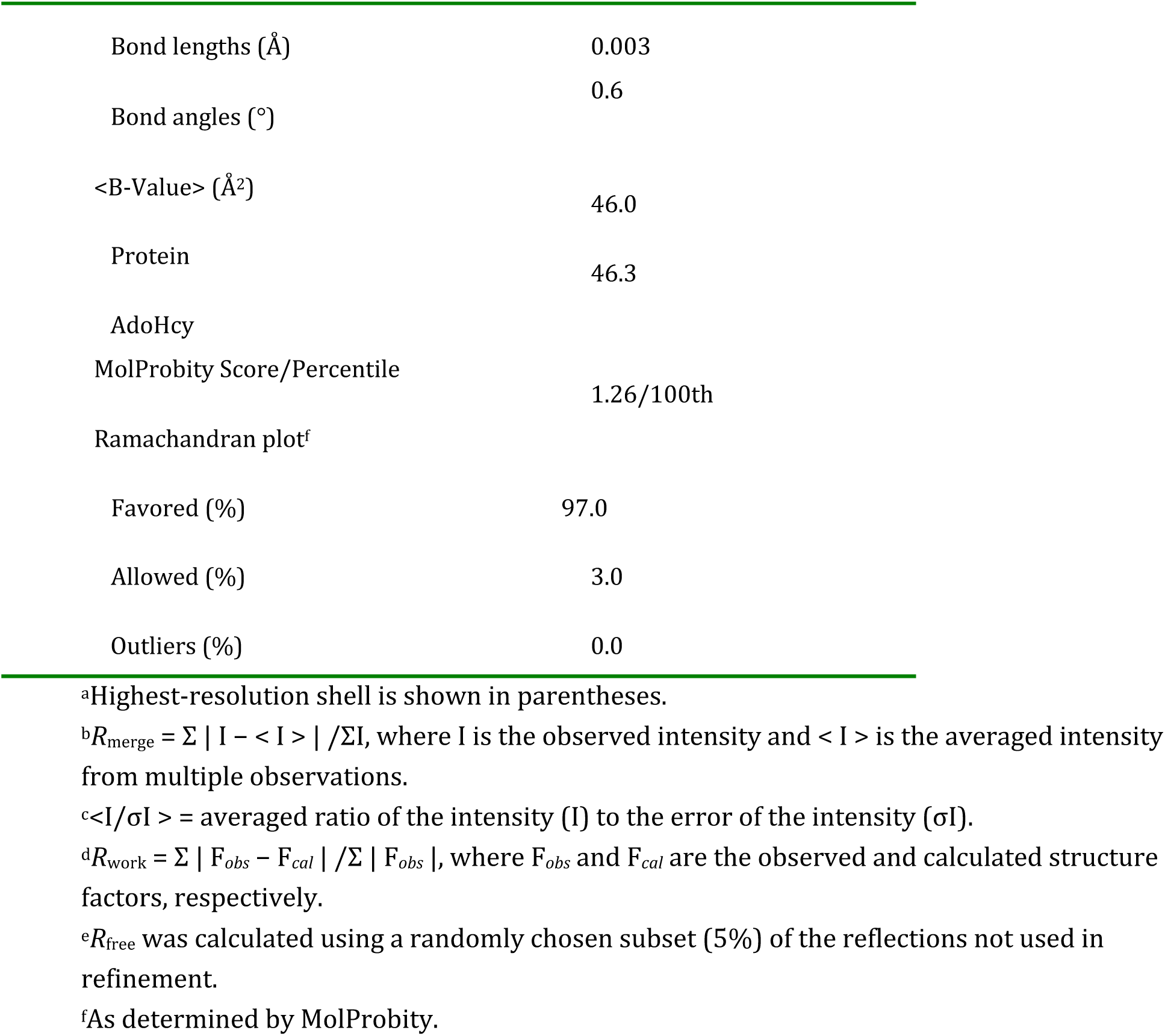
Data collection and refinement statistics.

|  |  |
| --- | --- |
| **Data collection** |  |
| Beamline | NE-CAT 24-ID-C (APS) |
| Space group | C2 |
| Cell dimensions |  |
| a, b, c (Å) | a=196.6, b=65.9, c=141.0 |
| $\alpha, \beta, \gamma$ (°) | $\alpha=\gamma=90, \beta=106.9$ |
| Wavelength (Å) | 0.9792 |
| Resolution (Å)<sup>a</sup> | 50.0-2.40 (2.49-2.40) |
| No. of unique reflections | 62,179 (3,760) |
| $R_{\text{merge}}$ (%)<sup>a,b</sup> | 7.4 (70.6) |
| CC1/2<sup>a</sup> | 0.997 (0.673) |
| CC*<sup>a</sup> | 0.999 (0.897) |
| $\langle I / \sigma I \rangle^b$ | 24.0 (1.6) |
| Completeness (%)<sup>a</sup> | 88.8 (54.0) |
| Redundancy<sup>b</sup> | 7.3 (3.4) |
| **Refinement** |  |
| Resolution (Å) | 50.0 – 2.40 |
| No. of reflections | 55,457 |
| Test set | 1999 |
| $R_{\text{work}}^d / R_{\text{free}}^e$ (%) | 19.0 / 22.3 |
| No. of atoms | 10,318 |
| R.m.s deviations |  |
| Bond lengths (Å) | 0.003 |
| Bond angles (°) | 0.6 |
| <B-Value> (Å<sup>2</sup>) | 46.0 |
| Protein | 46.3 |
| AdoHcy |  |
| MolProbity Score/Percentile | 1.26/100th |
| Ramachandran plot<sup>f</sup> |  |
| Favored (%) | 97.0 |
| Allowed (%) | 3.0 |
| Outliers (%) | 0.0 |

<sup>a</sup>Highest-resolution shell is shown in parentheses.
<sup>b</sup> $R_{\text{merge}} = \sum |I - \langle I \rangle| / \sum I$ , where $I$ is the observed intensity and $\langle I \rangle$ is the averaged intensity from multiple observations.
<sup>c</sup> $\langle I/\sigma I \rangle$ = averaged ratio of the intensity ( $I$ ) to the error of the intensity ( $\sigma I$ ).
<sup>d</sup> $R_{\text{work}} = \sum |F_{\text{obs}} - F_{\text{cal}}| / \sum |F_{\text{obs}}|$ , where $F_{\text{obs}}$ and $F_{\text{cal}}$ are the observed and calculated structure factors, respectively.
<sup>e</sup> $R_{\text{free}}$ was calculated using a randomly chosen subset (5%) of the reflections not used in refinement.
<sup>f</sup>As determined by MolProbity.

**Supplemental Table S2.**

| | PDB accession | Identical | rmsd | No. of Cα atoms |
| --- | --- | --- | --- | --- |
| *Rn*PRMT1<sup>a</sup> | **<u>1OR8</u>** | 51% | 1.0 | 312 |
| *Sc*RMT1<sup>b</sup> | **<u>1G6Q</u>** | 47% | 1.2 | 308 |
| *Rn*PRMT3<sup>a</sup> | **<u>1F3L</u>** | 41% | 1.3 | 308 |
| *Mm*CARM1<sup>c</sup> | **<u>2V74</u>** | 35% | 1.6 | 302 |
| *Tb*PRMT1-PRO<sup>d</sup> | **<u>6DNZ</u>** (This study) | 24% | 2.1 | 293 |

<sup>a</sup>*Rn*: *Rattus norvegicus* (rat)
<sup>b</sup>*Sc*: *Saccharomyces cerevisiae* (budding yeast)
<sup>c</sup>*Mm*: *Mus musculus* (mouse)
<sup>d</sup>*Tb*: *Trypanosoma brucei*

